# Interacting cells driving the evolution of multicellular life cycles

**DOI:** 10.1101/533745

**Authors:** Yuanxiao Gao, Arne Traulsen, Yuriy Pichugin

**Affiliations:** Max Planck Institute for Evolutionary Biology, August-Thienemann-Str. 2, 24306 Plön, Germany

## Abstract

Evolution of complex multicellular life begun from the emergence of life cycle involving formation of cell clusters. Opportunity for cells to interact within clusters provided them an advantage over unicellular life forms. However, what kind of interactions may lead to the evolution of multicellular life cycles? Here, we combine evolutionary game theory with a model for the emergence of multicellular groups to investigate how cell interactions can influence reproduction modes during the early stages of the evolution of multicellularity. We identify evolutionary optimal life cycles as those which maximize the population growth rate. Among all interactions captured by two-player games, only eight life cycles were found to be evolutionarily optimal. Moreover, the vast majority of games promotes either of two classes of life cycles: (i) splitting into unicellular propagules or (ii) fragmentation into two offspring clusters of equal (or almost equal) size. Our findings indicate that the three most important characteristics, determining whether multicellular life cycles will evolve, are average performance of homogeneous groups, heterogeneous groups, and solitary cells.

## 1 Introduction

The evolution of multicellular life cycles is one of the most challenging questions of modern evolutionary biology. In the history of life, multicellular organisms have independently originated at least 25 times from unicellular ancestors [Grosberg and Strassmann, 2007]. From the very beginning, multicellular life has been shaped by interactions between different cells within heterogeneous groups [Okasha, 2006, Godfrey-Smith, 2009]. The role of these interactions in the emergence (or prevention) of multicellularity is an open question. Recently, there is a rising interest in the evolution of life cycles including multicellular stages from both experimentalists [Rossetti et al., 2011, Ratcliff et al., 2012, 2013b,a, Hammerschmidt et al., 2014] and theoreticians [Rainey and Kerr, 2010, Tarnita et al., 2013, Libby et al., 2014, De Monte and Rainey, 2014, Rashidi et al., 2015, van Gestel and Tarnita, 2017, Pichugin et al., 2017]. In the present study, we focus on competition between various multicellular and unicellular life cycles. The life cycle that leads to the fastest population growth would eventually dominate the population. We address how interactions between different cells within heterogeneous groups affect the growth competition between unicellular and multicellular life cycles. When interactions between different types of individuals within one group accelerate growth, more complex forms of multicellularity are expected to evolve in the long run.

We design our study with two specific scenarios of interacting cells in mind: the threat of free-riders in groups relying on cooperation and division of labour between cells. The very first multicellular organisms are commonly suggested to be composed from similar cells as suggested by fossils [Knoll, 1992, Tomitani et al., 2006] and experimental studies [Ratcliff et al., 2012, 2013b,a]. Cooperation between cells in these early organisms provided them benefits unavailable to solitary cells. However, free-riders gaining the cooperation benefits without paying any costs have an evolutionary advantage over cooperators, which in turn may violate the integrity of an organism [Hardin, 1968, Bonner, 1959, Rainey and Rainey, 2003]. One of the most efficient ways of policing free-riders is reproduction via single cell bottleneck, where an organism grows from a single cell. This suggests that interactions between cooperators and free-riders promote group reproduction with unicellular propagules.

The second scenario where cell interactions might play a significant role emerges once undifferentiated multicellularity has been established and cells began to specialize in various tasks. For example, consider filamentous cyanobacteria. During nitrogen depletion, cells in the filaments occasionally differentiate into nitrogen-fixating heterocysts that obtain sugars from neighbouring photosynthetic cells and, in turn, provide these cells with nitrogen. These heterocysts suffer a significant penalty to their own fitness, but are essential to the survival of the colony as a whole [Flores and Herrero, 2010]. A group reproduction mode preserving the necessary association between photosynthetic and rare nitrogen fixating cells would contribute a lot to the sustainable growth of this species. Naturally, the reproduction of cyanobacteria occurs by fragmenting the parental filament into shorter multicellular chains through programmed cell death [Claessen et al., 2014], so newly emerged multicellular colonies likely contain heterocysts and benefit from the division of labour from the very beginning. This suggests that the division of labour promotes group reproduction modes with multicellular offspring groups.

While there is no clear experimental evidence that the evolution of reproduction modes can be influenced by the interaction between cells of different types, such a hypothesis deserves a close attention. There is a range of previous models investigating the evolution of the division of labour [Gavrilets, 2010, Ispolatov et al., 2012, Rodrigues et al., 2012, Cooper and West, 2018]. However, these models incorporate a single predetermined reproduction mode, or a small hand-picked set of these. The evolution of cooperation in early multicellu-larity gained more attention [Michod, 1997, Nowak, 2006a, van Veelen, 2009]. Given that reproduction via single cell bottlenecks is a natural policing mechanisms, some aspects of the evolution of reproduction modes have been considered before. Examples are the evolution of propagule size [Roze and Michod, 2001, Michod and Roze, 2001], as well as the comparison between formation of cell clusters and unicellular existence [Kaveh et al., 2016]. However, the spectrum of possible interactions between cells goes way beyond specific scenarios of cooperation and the division of labour, so this topic remains largely unexplored.

In our study, we utilize the framework developed in [Pichugin et al., 2017], in which a reproduction mode is considered as a way to partition the cells comprising the parent group into two or more offspring groups. Since there is always a finite number of cells in a reproducing group, there is a finite number of possibilities for group fragmentation. However, our previous study assumed homogeneous groups composed of a single cell type. Here, we investigate heterogeneous groups consisting of cells of two different types. To represent the wide spectrum of possible interactions between two types, we utilize a game theory approach, where the result of cell interactions are given by payoff values derived from the payoff matrix of a given game. The payoff values affect both the growth rate of the whole group as well as difference in growth rates of cells within group. The combination of the game played in a group with the fragmentation mode determines the population growth rate. By screening a wide range of fragmentation modes, we find the one providing the largest growth rate, which is considered to be evolutionarily optimal reproductive strategy for the given game, as it leads to the fastest growth of biomass. Interestingly, when group growth is independent of the group size, only eight life cycles can be evolutionarily optimal among all possible games.

## 2 Model

We consider a group-structured population, where individuals of two phenotypes *A* and *B* are nested into groups. These groups incrementally grow by one cell at a time and fragment into smaller offspring groups upon reaching a critical size of *M* cells. For a given group, the time between cell divisions depends only on the size of this group and its cell composition. Thus, the growth of the group is independent of others and therefore at the level of groups, the population growth is density independent. Therefore, in the long run, the population converges to a stationary regime, characterized by exponential growth at a rate we call *λ*. As populations employing different life cycles (different critical size and/or fragmentation mode) have different growth rates, the life cycle with the largest growth rate *λ* will eventually take over the population.

### 2.1 Cell payoff and cell division

Interactions among cells in a group are captured by a pairwise game. The game is determined by a 2 × 2 payoff matrix

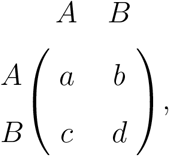

where *A* gets payoff *a* or b from interacting with *A* or *B* respectively, whereas *B* gets *c* or *d* from *A* or *B*, respectively. The average payoffs are given by

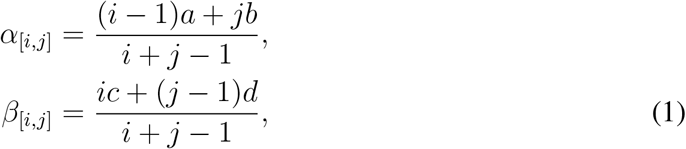

where *α*_[*i,j*]_ and *β*_[*i,j*]_ are the average payoff of *A* type cells and *B* type cells in a group of *i A*-cells and *j B*-cells, respectively (the −1 arises from the exclusion of self interactions, but such self interactions have only a minor influence on our results). Solitary cells do not play the game and their payoff is zero, so *α*_[1,0]_ = *β*_[0,1]_ = 0.

The role of a cell’s payoff is two-fold. First, cells with larger payoff have higher chance to reproduce, when the group grows incrementally. Second, groups with larger average payoff grow faster. We assume that selection is weak, which allows us to restrict ourselves to linear relationship between payoff of cells and their probabilities to divide as well as the time needed for the group to grow.

Once a cell division occurs, the probability of a cell to be chosen to divide increases linearly with its payoff, *P* ~ 1 + *wα* if the cell is of type *A*, and *P* ~ 1 + *wβ* if the cell is of type *B*, where *w* ≪ 1 is the selection strength. Therefore, the probabilities that the dividing cell will be of type *A* or *B* under weak selection, *w* ≪ 1, are

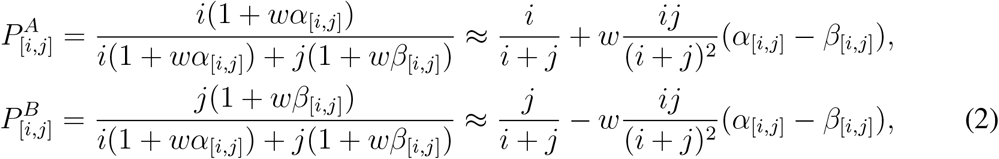

where 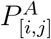 is the probability that some cell of type *A* will be chosen to divide in a group of *i A*-cells and *j B*-cells, and 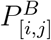 is the same for type *B*, so 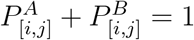.

Similarly, the time between two consecutive cell divisions depends linearly on the average payoff in a group

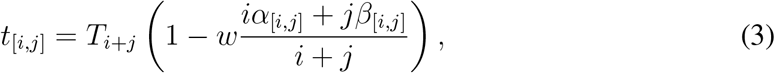

where *T*_*i*+*j*_ is the size dependent component of growth, and 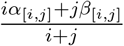 is an average payoff of cells in a group.

When a cell divides, each of the two daughter cells may independently change their type with probability *m*. Thus, the daughter cells of an *A* cell are either two *A*-cells with probability (1 − *m*)^2^, or one *A* and one *B* cell with probability 2*m*(1 − *m*), or two *B*-cells with probability *m*^2^.

Once the group reaches the critical group size *M*, it immediately fragments into smaller pieces and all cells are randomly assigned to offspring groups. The life cycle is determined by the critical size *M* and the sizes of offspring groups. For instance, at *M* = 3, there are two possible life cycles: either split into three solitary cells (life cycle 1+1+1), or into a solitary cell and bi-cellular group (life cycle 2+1). For *M* = 4, there are four possible life cycles: 3+1, 2+2, 2+1+1 and 1+1+1+1. Below, we refer to different life cycles using partitions of integer numbers.

### 2.2 Population growth rate

We assume that the population can grow without any bounds. For our model, the density of groups follows a linear differential equation and growth is exponential [Pichugin et al., 2017]. Our goal is here to find the overall population growth rate *λ*.

To do so, we need to take into account the stochastic nature of group development in our model. There are three sources of stochasticity: (i) the choice of the cell to divide, (ii) the phenotype of daughter cells born at the cell division, and (iii) the assignment of cells to offspring groups at group fragmentation. As a consequence, groups are born different: a newborn bi-cellular group may consist of two *A*-cells, one *A* cell and one *B* cell, or two *B*-cells. Also, due to the randomness in outcomes of individual cell divisions, initially identical groups may follow different developmental trajectories during their growth, where by “developmental trajectory”, we mean the record of all choices made among possible alternatives during the group growth.

Fortunately, the number of newborn states and the cell composition after each division is finite, see Fig. 1. Therefore, for any life cycle, we take all possible developmental trajectories into account. For an arbitrary life cycle, each group is born as one of S initial types, which we enumerate as (1, 2, ⋯, *S*). For each available developmental trajectory t, we designate the initial state of the trajectory as *i*(*τ*), the probability that a group born at initial state *k* will follow the trajectory as *p_k_*(*τ*), such as *p_k_*(*τ*) = 0, if *k* ≠ *i*(*τ*), the time necessary to complete the trajectory as *T*(*τ*), and the vector of numbers of each offspring type produced at the end of the trajectory as **N**(*τ*) = (*N*_1_, *N*_2_, ⋯, *N_S_*).

**Figure 1:**
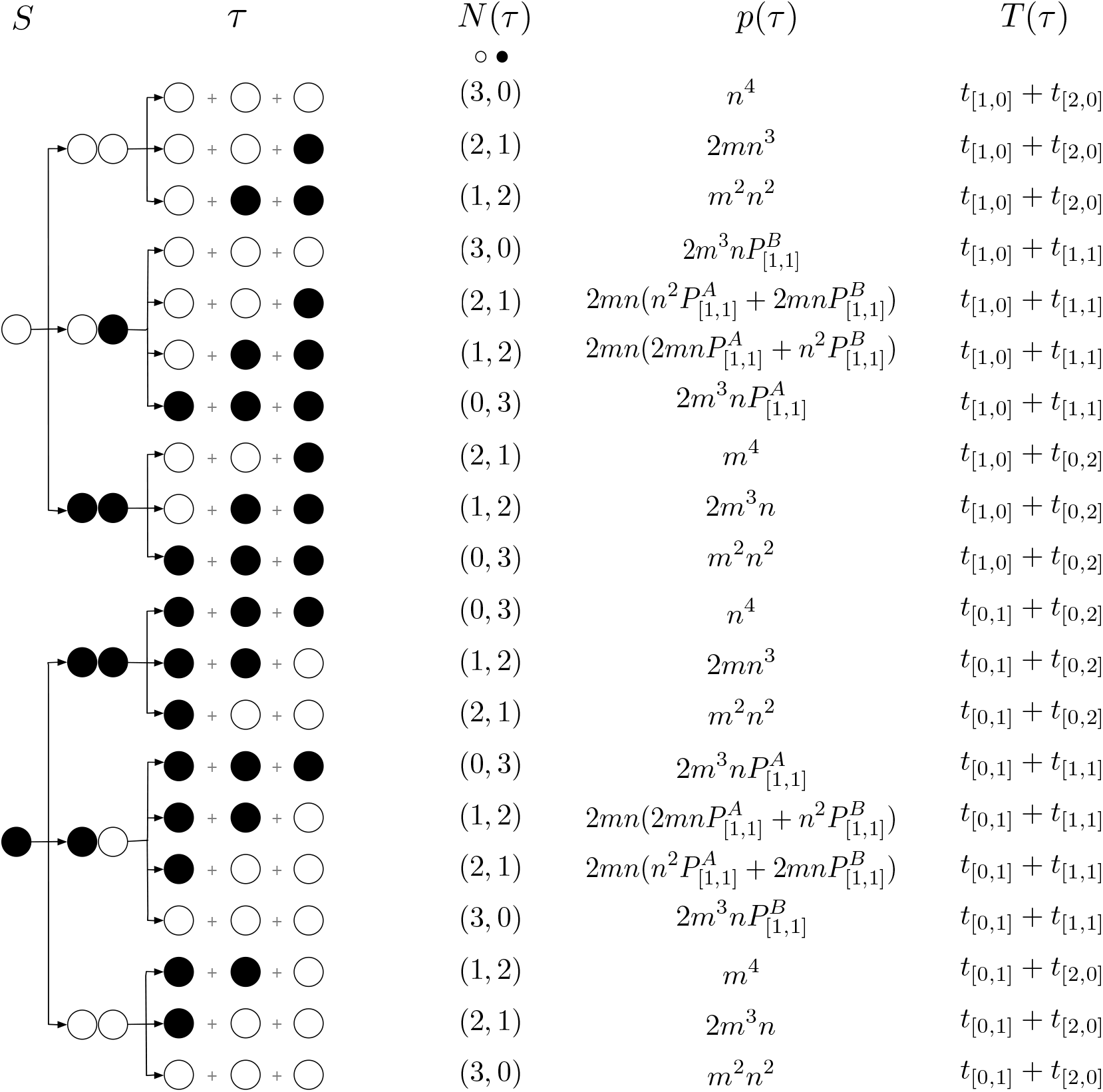
The number of developmental trajectories is finite. Here we show the full set of developmental trajectories (*τ*) in the life cycle 1+1+1, where groups are born unicellular, then grow up to size three and immediately split into independent cells. This life cycle features only two initial states *S*: solitary *A*-cell (open circles) and solitary *B*-cell (black circles). Stochastic phenotype switching creates 10 possible developmental trajectories for each initial state. To shorten the notation, we use *n* = 1 − *m* - the probability of a daughter cell to have the same phenotype as the mother cell.

The growth rate of population *λ* is given by the solution of equation (Appendix A.1)

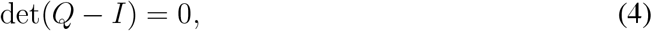

where *I* is the identity matrix and *Q* is a matrix, in which

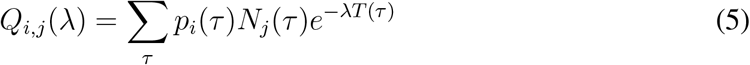

is the contribution of groups born as type *i* to the production of newborn groups of type *j*, see also [De Roos, 2008].

## 3 Results

Our model allows to calculate the growth rate of any given life cycle provided the elements of the payoff matrix (*a, b, c, d*), the phenotype switching probability *m*, and the profile of size-dependent component of development time (*T*_*i*+*j*_). Here, we focus on the life cycles having the largest λ, as these will be the winners of evolutionary growth competition.

In our study, we assume that in the absence of interactions (*w* = 0), all cells divide independently at the same rate. Consequently, the time of doubling of the group size is the same for groups of any size. This implies 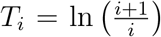 and as we show in Appendix A.2, at *w* = 0 this leads to the same population growth rate *λ* = 1 for all life cycles. We also considered other developmental times profiles at *w* = 0 and the results of our model are similar to our previous investigation of life cycles of homogeneous groups [Pichugin et al., 2017], see Appendix A.3.

Under weak selection, the growth rate of population with an arbitrary life cycle *κ* can be approximated by 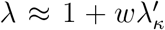. The expression for 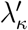 can obtained from the linearisation of Eq. (4) with respect to *w*. Since the payoffs *a, b, c, d* always come into play with a factor *w* (see Eqs. (2), (3)), 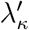 is also linear in these payoffs. The dynamics of the population as a whole does not change if we exchange the two cell types *A* ↔ *B* and the corresponding payoff values *a* ↔ *d, b* ↔ *c*. Thus, *a* and *d* contribute to 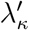 with the same weight and the same is true for *b* and *c*. Therefore, 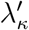 can be presented as a function of only three parameters: *m*, *ψ* = *a* + *d* and 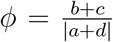. The parameter *ψ* can be interpreted as whether the formation of a homogeneous group is beneficial to the cell (*ψ* > 0) or not (*ψ* < 0), compared against the case of the solitary cell. The value of *ϕ* can be interpreted as the benefit from the formation of a heterogeneous group compared to the formation of a homogeneous group. In a broad sense, *ϕ* captures how well groups of mixed composition perform against pure groups. The details of the calculation of 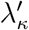 can be found in Appendix A.4.

We numerically investigated the optimality of life cycles with fragmentation size *M* up to 7. In total, there are 37 such life cycles, see Fig. 2. However, only eight of them were found to be evolutionarily optimal for any combination of control parameters, see Fig. 3. These life cycles fall into one of three categories: fission into multiple unicellular offspring (1+1, 1+1+1, 1+1+1+1, and 1+1+1+1+1); binary fragmentation with group propagules (2+2 and 4+3); and the rarely observed transition between the previous two classes (2+1 and 2+1+1).

**Figure 2:**
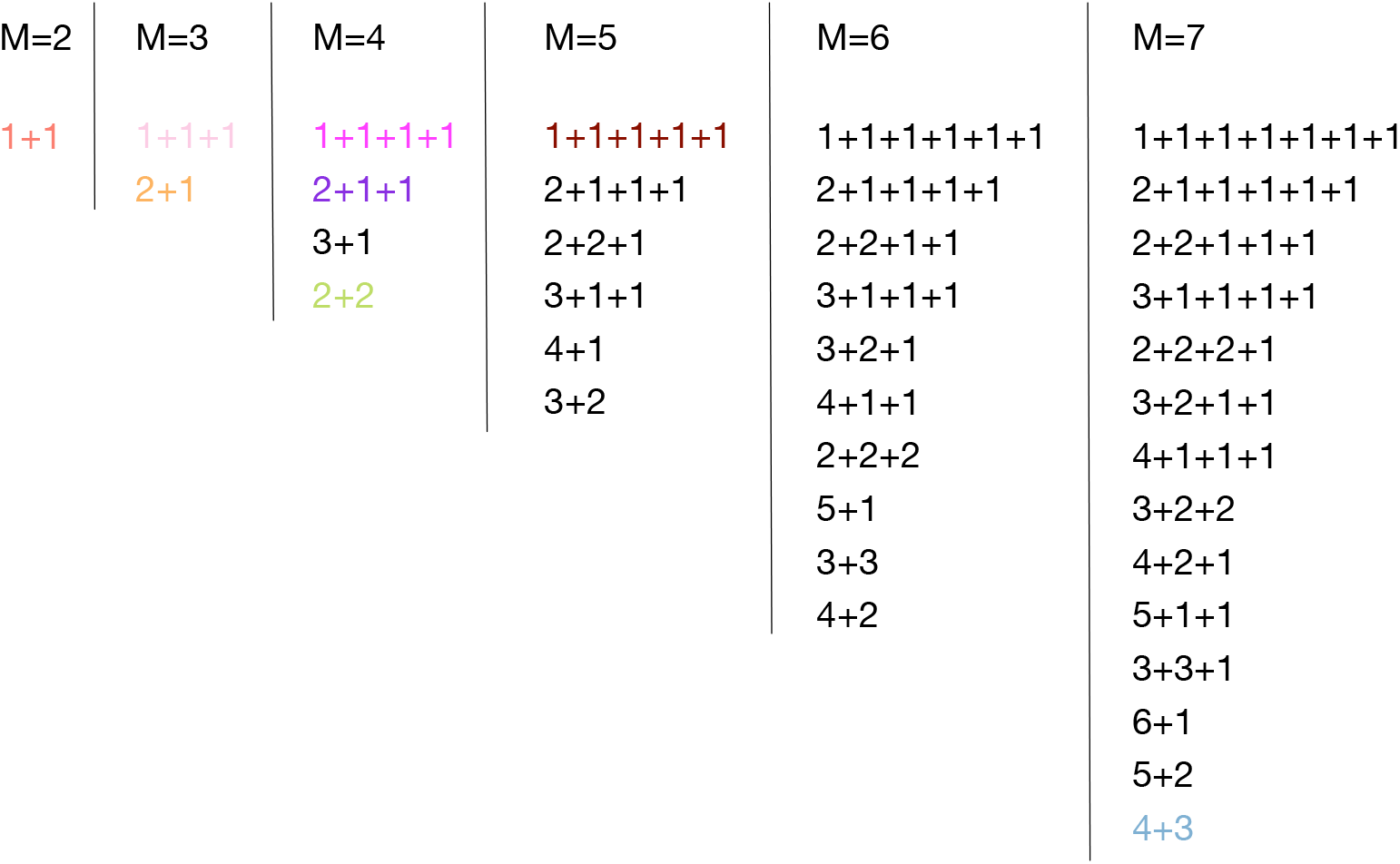
The list of all life cycles with critical sizes. *M* ≤ 7. The coloured life cycles are those found to be evolutionarily optimal for some combination of the control parameters *m, ψ* and *ϕ*. Most of life cycles were never found to be optimal. Among 24 life cycles corresponding to two largest critical sizes *M* = 6 and *M* = 7, only one is found to be evolutionary optimal – 4+3.

**Figure 3:**
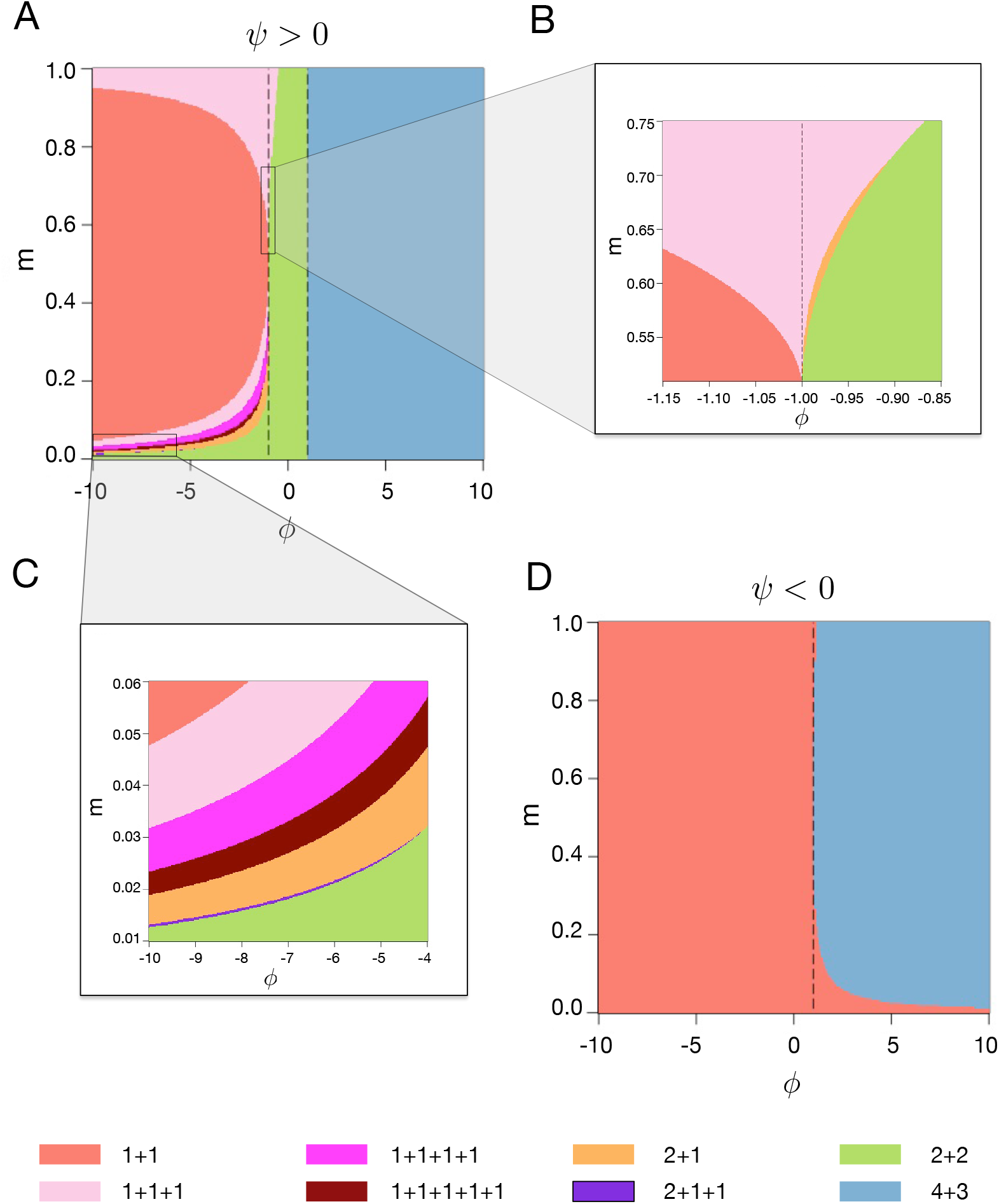
Only eight life cycles are evolutionarily optimal under weak selection. Panel A: Optimal life cycles for *ψ* > 0. Dashed lines are *ϕ* = −1 and *ϕ* = 1. Panel B: The transition under large phenotype switching rate *m* > 0.5 and *ψ* > 0. Panel C: The transition under small phenotype switching rate *m* ≪ 1 and *ψ* > 0. Panel D: Optimal life cycles under *ψ* < 0. Dashed line is *ϕ* = 1.

Among these eight, only the life cycle 4+3 reflects the imposed limit of the maximal group size equal to seven cells. We observed that if the group size is limited to *M* ≤ 5, the life cycle 3+2 can be evolutionary optimal. Extending the size limit to *M* ≤ 6, that life cycle is replaced by 3+3, and finally at *M* ≤ 7, the life cycle 4+3 takes this place. Therefore, the life cycle 4+3 listed in this study is likely the manifestation of the more general rule “grow as large as possible and divide into two equal or almost equal parts”.

We break the analysis of our results into two parts. First, we consider specific life cycles and outline the conditions which promote their evolution. Then, we take the opposite direction and focus on specific games to investigate which life cycles are promoted by them.

### 3.1 Games promoting a given life cycle

First, we examine the optimal life cycles for negative *a* + *d* (*ψ* < 0), in which homogeneous groups are in adverse conditions in the first place, see Fig. 3D. Consequently, one of two life cycles found here is 1+1 – unicellularity, at which groups are not formed at all. Still, if ϕ is sufficiently large, the highest growth rate is obtained by heterogeneous groups. Then, evolutionary growth competition favours life cycles minimizing the fraction of homogeneous groups in the population. Due to the random partitioning of cells into offspring groups, smaller offspring have larger chances to accumulate cells of only one type during fragmentation. Thus, growth competition would likely promote larger offspring size to avoid such outcomes. If so, the optimal life cycle must be the fragmentation into two equal-sized (or nearly equal) offspring group at the maximal available size (4+3 in our case). Next, we focus on the more complex case of *ψ* > 0, see Fig. 3A.

When *ϕ* > 1, the life cycle 4+3 is evolutionarily optimal. At these values of *ϕ*, all groups have an advantage over solitary cells, but heterogeneous groups profit more than homogeneous. Therefore, growth competition favours life cycles avoiding production of independent cells and minimizing the fraction of homogeneous groups in the population, i.e. equal binary split at the maximal size. Note that at *ϕ* = 1, where *a* + *d* = *b* + *c*, there are no benefit differences between homogeneous and heterogeneous groups. As a consequence, all life cycles with multicellular offspring have the same growth rate there, see Fig. 4B.

**Figure 4:**
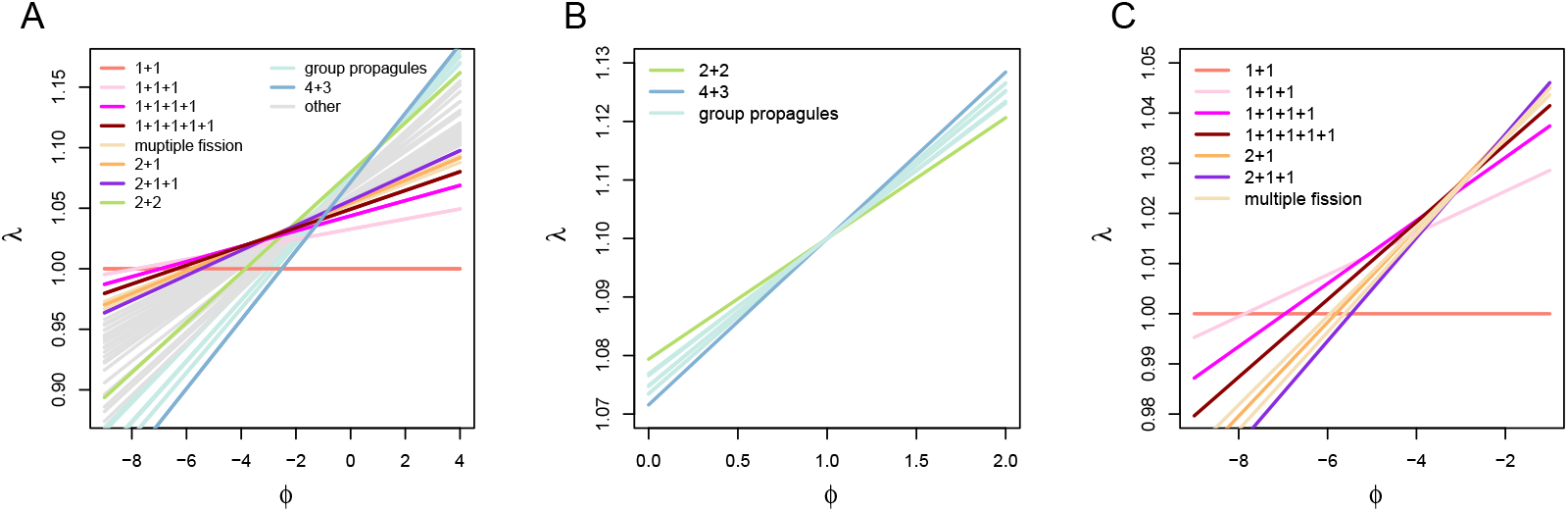
The growth rates of the considered life cycles as a function of *ϕ* for *ψ* > 0. Panel A: according to the weak selection approximation, growth rates *λ* are linear functions of *ϕ*. For all life cycles, the slope of the line is non-negative, thus, life cycles with smaller slope dominate at *ϕ* ≪ −1 (1+1 has slope zero) and life cycles with larger slope dominate at *ϕ* » 1 (4+3 has the largest slope for *M* ≤ 7). Panel B: all life cycles with multicellular offspring share the same growth rate at *ϕ* = 1 (*ϕ* = −1 under *ψ* < 0). Panel C: a sequence of multiple fission life cycles is optimal at the negative *ϕ*. At all panels *m* = 0.06. In all panels, multiple fission includes 1+1+1+1+1+1, 1+1+1+1+1+1+1; group fission includes 3+2, 3+3, 2+2+2, 4+2, 5+2. Growth rates profiles at other parameters are presented in Appendix A.5.

For 0 < *ϕ* < 1, 2+2 is the optimal life cycle. Here, all groups have an advantage over solitary cells but homogeneous groups benefit more than heterogeneous. Therefore, growth competition would likely to promote life cycles maximising the fraction of homogeneous groups in a population. First, this means producing the smallest multicellular offspring (bi-cellular groups) to eliminate parental heterogeneity in offspring. Second, the fragmentation has to be performed at the smallest size to minimize the risk of gaining heterogeneity in groups due to a spontaneous phenotype switch during growth. For the bi-cellular offspring, the smallest fragmentation size is four cells, therefore, the best life cycle must be 2+2. Interestingly, if *m* is small enough, 2+2 life cycle can be optimal under arbitrary large negative *ϕ*, see Fig. 3A, C. There, while heterogeneous groups have a strong disadvantage, chances of the phenotype switch to occur are low and homogeneity of groups is generally preserved.

At *ϕ* < 0, the emergence of another cell type in homogeneous groups incurs a penalty on the group growth. To avoid the production of heterogeneous groups, growth competition is likely to promote life cycles involving dispersal into independent cells, such that each newborn group starts in a homogeneous state.

When *ϕ* < 0 and *m* is high enough, heterogeneous groups are likely to form after the very first cell division. In this case, 1+1 is favoured as it does not involve any group formation at all. However, once *m* approaches zero, the first few cell divisions performed by initially solitary cell will likely produce a homogeneous group. Thus, multicellular life cycles with fission into independent cells are favoured: 1+1+1, 1+1+1+1, and 1+1+1+1+1,see Figs. 3C and 4C. Larger fragmentation sizes: 3, 4, and then 5, become optimal with decreasing m. However, fission at size 6 has never found to be optimal, because at this stage, the production of multi-cellular offspring become beneficial despite the risk of transferring parent heterogeneity into the next generation.

Transitional life cycles 2+1 and 2+1+1 are found to be optimal between areas of optimality of multiple fission life cycles (1+…+1) and multicellular offspring life cycles (2+2 and 4+3), see Fig. 3C. These two life cycles mix unicellular and multicellular offspring. This may be a result of compromise between producing multicellular offspring to fully utilize benefits of interactions in homogeneous groups, and the necessity to fragment into independent cells to purge emerging heterogeneous groups.

### 3.2 Life cycles promoted by prominent games

The most prominent game in the context of evolutionary game theory is the Prisoner’s dilemma [Weibull, 1995, Nowak, 2006a, Pacheco et al., 2009, Hilbe et al., 2013]. In the simplest form of the Prisoner’s dilemma, each player may pay some cost 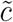, so that the opposing player will receive a benefit 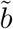 (larger than the cost). The cooperating strategy is to pay the cost, while the defecting strategy is to abstain from paying this cost (but still receive incoming benefits). The largest combined payoff is achieved by both players cooperating, while the individual’s payoff resulting from defecting behaviour is always larger than payoff from mutual cooperation. The conflict between individual’s and group’s interests makes this game a social dilemma.

The payoff matrix of the simplest Prisoner’s dilemma is given by

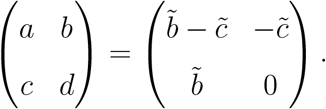

With these payoffs, 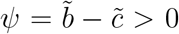 and 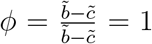. Surprisingly, this game exhibits a special behaviour in our model: any life cycle which does not pass through unicellular stage (e.g. 3+2+2) is evolutionarily optimal, independently of the phenotype switch probability *m* (i.e. risk of defector emergence). Contrary to intuition, cooperative cell interactions described by the Prisoner’s dilemma promote everything *except* the reproduction via single cell bottleneck. This is due to the fact that in a group with at least one cooperator, some benefit is already produced and shared across the group. Thus, preserving group living is more advantageous for the population than producing single cell propagules.

Other notable social dilemmas are snowdrift and stag hunt games. In the snowdrift game, a combined cost 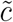 must be paid for the benefit 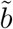 to be received by each player. Cooperators readily pay their share of the costs, while defectors abstain from paying it. The payoff matrix of the snowdrift game is

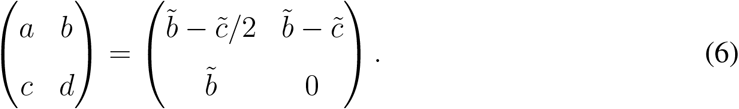

This results in 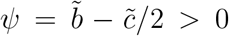 and 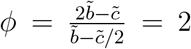. According to our findings, these parameters promote the life cycle 4+3, or, more generally, equal binary fragmentation at the maximal possible size, which ensure heterogeneous groups that maximize the combined payoff.

In the stag hunt game, players may pursue a hare – small prey providing payoff *h*, or a stag - large prey giving payoff *s* > *h*. Hare hunt is always successful, but only both hunters together can hunt down the stag. The payoff matrix of the stag hunt game is

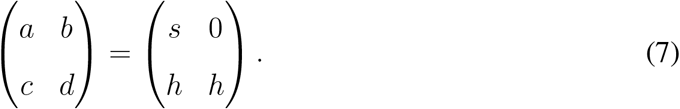

This results in *ψ* = *s* + *h* > 0 and 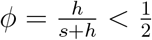. These parameters promote the life cycle 2+2. In contrast to the Prisoner’s dilemma and snowdrift games, in the stag hunt game a group of mixed composition has the smallest combined payoff (which is still larger than zero payoff for solitary cells). Therefore, the stag hunt game strongly favours a life cycle preserving homogeneity of groups, i.e. 2+2.

Many other evolutionary games have been studied and applied in a wide variety of biological situations [Maynard Smith, 1982, Hofbauer and Sigmund, 1998, Nowak and Sigmund, 2004]. For the general case, in a large well-mixed population of players, three classes of evolutionary dynamics are possible in 2 × 2 games: dominance of one strategy (*a* > *c, b* > *d* or *a* < *c, b* < *d*), bistability (*a* > *c, b* < *d*) as well as coexistence (*a* < *c, b* > *d*) [Weibull, 1995, Nowak, 2006b].

All games experiencing a bistability (such as stag hunt) have *ϕ* < sign(*ψ*). According to our results, for positive *ψ*, bistability games can promote 7 out of 8 found life cycles: equal binary split at the maximal size (4+3) does never lead to the fastest growth rate. For negative *ψ* bistability games only lead to unicellular life cycle (1+1). Games featuring coexistence dynamics (such as snowdrift) satisfy *ϕ* > sign(*ψ*), which restricts the optimal life cycle to 4+3 under *ψ* > 0 but allows both 1+1 and 4+3 under *ψ* < 0. Dominance games (such as the Prisoner’s dilemma) may have any value *ϕ*, so they can promote any of 8 found life cycles.

## 4 Discussion

In our study we performed an extensive investigation of the competition of life cycles driven by interactions between cells within in a group. Key to this study is the consideration of all possible reproduction modes and all possible interactions captured by game theoretic 2 × 2 payoff matrices. Among the huge variety of reproduction modes, only eight were found to be evolutionarily optimal, see Figs. 2 and 3. Moreover, the vast majority of games promotes either of two very specific classes of life cycles: fragmentation into strictly unicellular offspring (1+…+1) or production of exactly two strictly multicellular daughter groups of identical (or almost identical) size. Intuitively, life cycles with unicellular offspring should be promoted when the cell grow fastest in a homogeneous group, as the single cell bottleneck eliminates heterogeneity in the most effective way. Similarly, when the cell grow fastest in a heterogeneous group, life cycles with multicellular offspring should be promoted as they are best in preserving heterogeneity. Our results, in general, support this intuition. However, the current work reveals a much broader picture and we observed a number of less intuitive features of life cycle evolution driven by cell interactions. First, we observed the transition between these two major life cycles classes. This occurs via transitional life cycles mixing unicellular and multicellular offspring (such as 2+1 and 2+1+1), see Fig. 3C. Second, we found that if being in a heterogeneous groups incurs a moderate penalty onto the cell, growth competition may still promote the life cycle with only multicellular offspring (2+2) even at high rates of phenotype switch (*m*), see Fig. 3A. Third, an arbitrary strong penalty to heterogeneous groups (*ϕ* < 0), may still lead to the evolution of life cycles with multicellular offspring (2+2) given small enough *m*, see Fig. 3A, C. Altogether, even with only eight life cycles observed, our model exhibits a rich behaviour and gives valuable insights into factors shaping the evolution of life cycles.

We found that social dilemma games may not promote the evolution of single cell bottlenecks. A naive intuition suggest life cycles with unicellular offspring to be favoured by all social dilemmas as single cell bottleneck is such an effective way to police defectors. However, social dilemmas may lead to the evolution of any of the eight life cycles. What would be the reason for such a counter-intuitive outcome?

A key difference between our approach and the most of studies utilizing evolutionary game theory is that while we allow the competition between different cell types (by means of different division probabilities *P^A^* and *P*^B^), winning in such a competition is not in the focus of our attention. We consider both cell types as essential components of the group development. This is in line with the previous idea of Rainey and Kerr [2010] that cheaters may play a significant role in the evolution of life cycles in early multicellularity. Embracing this approach, we acknowledge that life cycles showing the largest population growth rates are not necessarily the best in keeping cheaters out. Our results shows that for evolution to favour single cell bottlenecks, a group mixing cooperators and defectors should have lower average fitness than an equivalent pack of independent cells. Otherwise, life cycles with multicellular offspring will be promoted.

This lead us to the second key feature of our model: the role of solitary cells. Independent cells stand out as they have no other cell to interact with and, thus, do not play a game. As such, they serve as a benchmark of the cell behaviour, against which all other group compositions are compared. Our results indicate that optimality of life cycles strongly depends on whether a (homogeneous) group formation is beneficial or deleterious compared to a solitary cell, see Fig. 3 A and D, respectively. For the Prisoner’s dilemma game, a combination of a single cooperator and single defector, indeed harm the cooperator the most. However, the overall payoff to the group 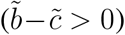 is still larger than the zero cumulative payoff these cells would obtain if separated. Thus, the Prisoner’s dilemma promotes the production of the multicellular offspring. The opportunity to abstain from the game (loner strategy)[Hauert et al., 2002, Fowler, 2005, Brandt et al., 2006, Hauert et al., 2007, Traulsen et al., 2009, García and Traulsen, 2012] is often viewed as component of the secondary importance in evolutionary game theory models, despite its potential impact on microbial dynamics [Garcia et al., 2014, 2015]. For the evolution of life cycles, such an opportunity plays a central role. For any life cycle producing unicellular offspring, each member of the population passes through a developmental stage without any interaction. Also, an ultimate loner strategy, where no game is ever played, is implemented by the unicellular life cycle, which is the most basic and one of the most important reproduction modes.

The interplay between cell interactions and life cycles has been considered in previous studies. Roze and Michod [2001] compared the growth rate of two reproductive modes: a spore reproducer (multiple fission life cycles in our terms) and the fragmentation into same sized offspring groups. Based on the fitness effects from the colony size, they investigated the situation which life cycle is good at eliminating the mutation (both selfish and deleterious mutation).

An explicit connection between fragmentation modes and games played within the group has been first made by [Kaveh et al., 2016]. There, authors focused on fragmentation modes in a form *x* + 1, and explicitly considered the 2+1 life cycle. Being focused on cooperation rather than evolution of life cycles, they dicsussed conditions promoting the evolution of cooperation.

The results of our model can be directly compared with our previous findings in [Pichugin et al., 2017] and [Pichugin and Traulsen, 2018], which considered the evolution of life cycles in homogeneous groups. There, for the costless fragmentation, as in the present study, only binary fragmentation modes (i.e. in a form *x* + *y*) can be evolutionarily optimal. Once the reproduction incurs a cost, fragmentation into multiple parts may evolve, still some fragmentation modes remain “forbidden”, i.e. they cannot evolve under any fitness landscape (*T_i_* in our terms). The set of evolutionarily optimal life cycles found in the current study is significantly different from the sets described above. Fragmentation in our model is costless, and yet we found that fragmentation into multiple parts may evolve due to the impact of cell interactions. Also, the life cycle 2+1+1, which may evolve in our model, belongs to the class of “forbidden” life cycles under the costly reproduction, so it cannot evolve among homogeneous groups at all. Thus, the introduction of a heterogeneity and interactions between different cell types make it possible for previously unattainable life cycles to evolve.

It is a challenging question how the interactions between different cells within an organism shape its reproduction mode. The present study demonstrates that this topic can be addressed systematically. To do so, we combine evolutionary game theory with the theory of life cycles in simple multicellular organisms. Game theory is capable to capture arbitrary interactions by a payoff matrix. At the same time, the theory of life cycles represents an arbitrary reproduction mode by the partition of an integer number. These two general frameworks naturally complement each other and allow holistic investigation of life cycles of organisms with heterogeneous composition, where it is impossible to evaluate the evolution of one factor neglecting another.

## A Appendix

### A.1 Population growth rate in the case of stochastic developmental programs

Consider a population in which each group emerges as one of *S* initial types. These types could be the newborn groups of different size and/or composition. With time passing, a group grows from its initial size to maturity and subsequent fragmentation. The set of growth events (cells divisions, mutations, etc) may vary from group to group. We call such an event chain “developmental trajectory” and designate it as *τ*. Any two groups of the same initial type may adopt different developmental trajectories for a number of reasons, such as mutations, stochastic developmental programs, or different environmental conditions. We use the following parameters of the developmental trajectory: *i*(*τ*) – the initial state of the group leading to the given developmental trajectory, *p_k_*(*τ*) – the probability that a group that emerged as initial type *k* will follow the trajectory *τ*, so *p_k_*(*τ*) = 0, if *k* ≠ *i*(*τ*), *T*(*τ*) – the time necessary to the newborn group to complete the trajectory *τ* and **N**(*τ*) = (*N*_1_, *N*_2_, ⋯, *N_S_*) – the vector of numbers of each offspring type produced during the fragmentation at the end of the trajectory *τ*.

The population features an explicit maturation component: a newborn group does not reproduce until time *T*(*τ*) has passed. Thus, to describe the population dynamics and find the population growth rate *λ*, it is necessary to consider the population demography. To do so, we characterize each group at each moment of time by the age parameter *η*. We define the age in a way that the newborn group has *η* = 0, while the group that reached the end of the developmental trajectory and is about to fragment has *η* =1. Along the trajectory, the age increases at a constant rate equal to 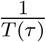, i.e. the rate of ageing differs between different trajectories.

From the perspective of the population dynamics, any two groups sharing the same developmental trajectory *τ* and age *η* are identical. Thus, the state of the whole population can be described by the density function *ζ* (*τ, η, t*), which shows how many groups on the developmental trajectory *τ* have age *η* at the given time *t*. In the stationary regime, where the fraction of groups of each type stays constant, the density function grows exponentially,

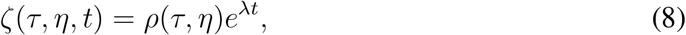

where *ρ*(*τ, η*) is the stationary density distribution of groups in a population.

Within a given developmental trajectory, ageing occurs at the same rate for all groups. Therefore, the dynamics of the density function at a given age *η* is determined by the balance between influx of maturing younger groups and the outflux of groups becoming too old. Both processes occur with the same rate 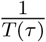, thus the density function must satisfy the transport equation

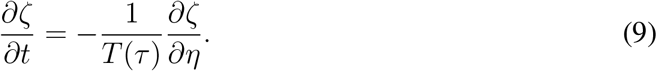

Combining Eqs. (8) and (9) we get

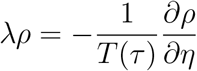

The solution of this equation is

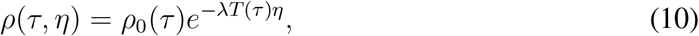

where *ρ*_0_(*τ*) is the stationary density distribution of newborn groups with *η* = 0.

To find *ρ*_0_(*τ*), we use the fact that each newborn organism is produced as a result of the fragmentation of some mature organism. Thus, the rate of emergence of newborn organisms in the population (*j*_0_) is the same as the rate of production of offspring in the course of reproduction of mature organisms (*j*_1_).

For any developmental trajectory *τ*, the rate of entering into the newborn state per time unit is equal to

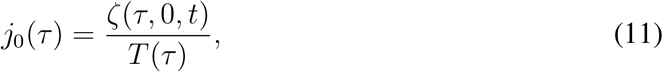

where the right hand side of the equation is the product of the number of newborn groups and the rate of ageing. The number of offspring with developmental trajectory *τ* is equal to the product of the total number of offspring of type *i*(*τ*) produced by all mature organisms and the probability of the offspring to adopt this developmental trajectory (*p_i_*(*τ*)(*τ*))

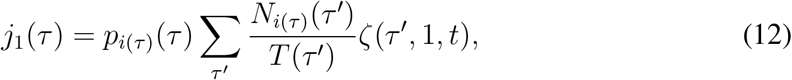

where summation is performed over all possible developmental trajectories of parent groups. Since each produced propagule is a newborn organism, *j*_0_(*τ*) = *j*_1_(*τ*). Therefore,

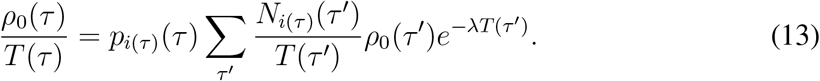

To obtain the expression connecting the population growth rate *λ* with parameters of developmental trajectories *τ*, we multiply both parts by *N_j_*(*τ*)*e*^−*λT*(*τ*)^ (note that in general *j* ≠ *i*(*τ*)) and sum over all possible developmental trajectories

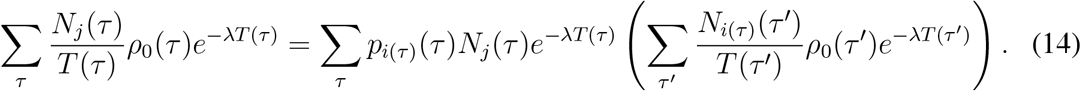

We define

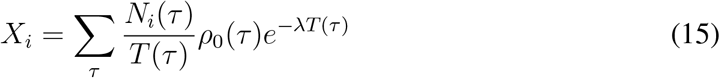

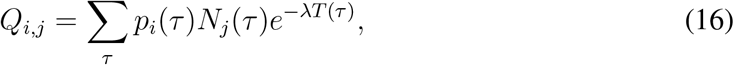

Note that *p_j_*(*τ*) = 0 if *j* ≠ *i*(*τ*).

Taking into account that *p_j_*(*τ*) = 0 if *j* ≠ *i*(*τ*), Eq. (14) becomes

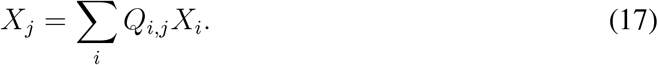

Also in the definition of *Q_i,j_*, the result of summation over all trajectories *τ* is the same as over only developmental trajectories starting from the initial state of type *j*, since *p_j_*(*τ*) = 0, if *j* ≠ *i*(*τ*), because an organism emerged as one type has no access to developmental trajectories originated from other types.

Eq. (17) can be satisfied only if

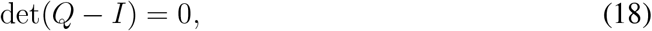

where elements of matrix *Q* are defined by Eq. (16) and *I* is identity matrix. This equation allows to infer the population growth rate *λ* if the parameters of each trajectory are known (*i*(*τ*), *p_i_*(*τ*), **N**(*τ*) and *T*(*τ*)). In most interesting cases, this has to be done numerically.

### A.2 Existence of the neutral fitness landscape in the case of homogeneous groups

Consider the situation, where *w* = 0 and, therefore, the group properties depend only on the group size. A group of size i grows in size to *i* + 1 within time *T_i_*. Here we show that if 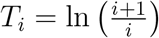, all life cycles have the same growth rate *λ* =1. We prove this by induction:

- The base of induction is given by Eq. (4), which states that if 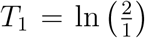 and 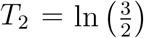, then *λ* =1 for any life cycles fragmenting at size 3 or smaller.
- The step of induction must show that if the assumption of induction holds true for maximal size *M*, then under adding 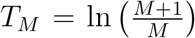, the assumption also holds true for maximal size *M* + 1. To prove the step of induction, we only need to consider life cycles fragmenting exactly at the size *M* + 1 because life cycles fragmenting at sizes smaller than *M* +1 have *λ* =1 according to the assumption of induction.

To construct the matrix *Q* and find the growth rate of considered life cycles, we need to characterize the set of offspring and developmental trajectories. In an arbitrary life cycle, the fragmentation of a homogeneous group of size *M* results in production of offspring groups of sizes ranging from 1 to *M*. In total, *M* different types of offspring can be produced, so the size of the matrix *Q* is *M* by *M*. Each of the offspring will grow up to size *M* + 1 and then fragment, thus there is only one developmental trajectory for each type of offspring with *p_i_*(*τ*) = 1. The developmental time of the trajectory 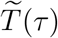 is given as the sum of incremental growth time

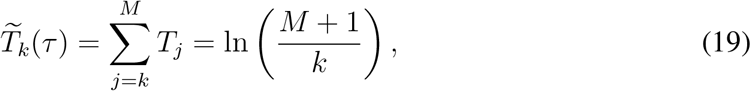

where *k* denotes the size of the newborn offspring.

An arbitrary life cycle can be characterized by the distribution of offspring sizes produced upon fragmentation *N_i_*, where *i* denotes the size of offspring. By the conservation of cell number during reproduction 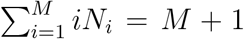. Therefore, according to Eq. (16), for an arbitrary life cycle, the elements of matrix *Q_ij_* are given by

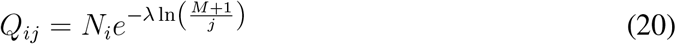

To prove the step of induction, we verify whether *λ* =1 is the solution of Eq. (18), with matrix *Q* given by Eq. (20). Plugging *λ* =1 into Eq. (20), we have 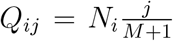, so the Eq. (18) becomes

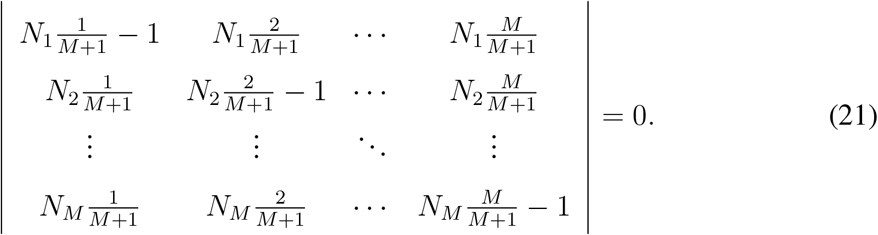

Based on the properties of determinant, we can take out the coefficients of each row and each column, then the left hand side of Eq. (21) becomes

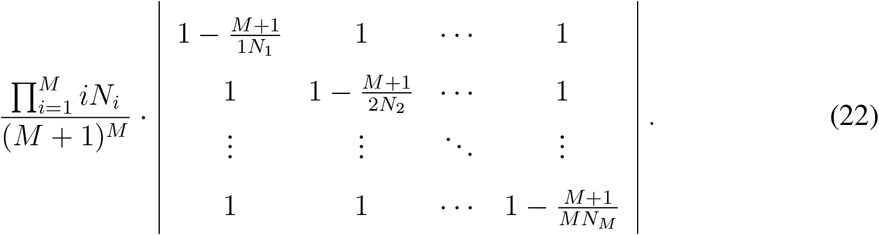

For convenience, we neglect the coefficient and denote 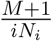 as *K_i_*. Thus, the determinant is

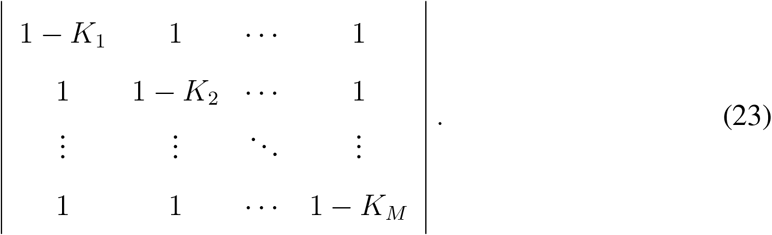

Next we calculate the determinant by splitting the first row,

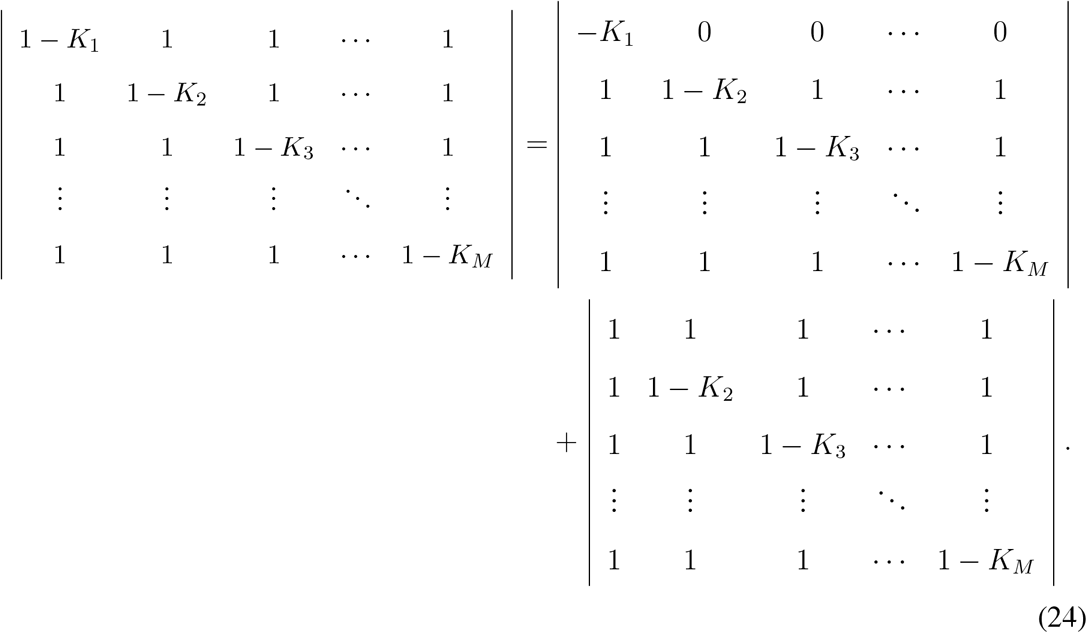

For the second part, splitting the second row, we can get

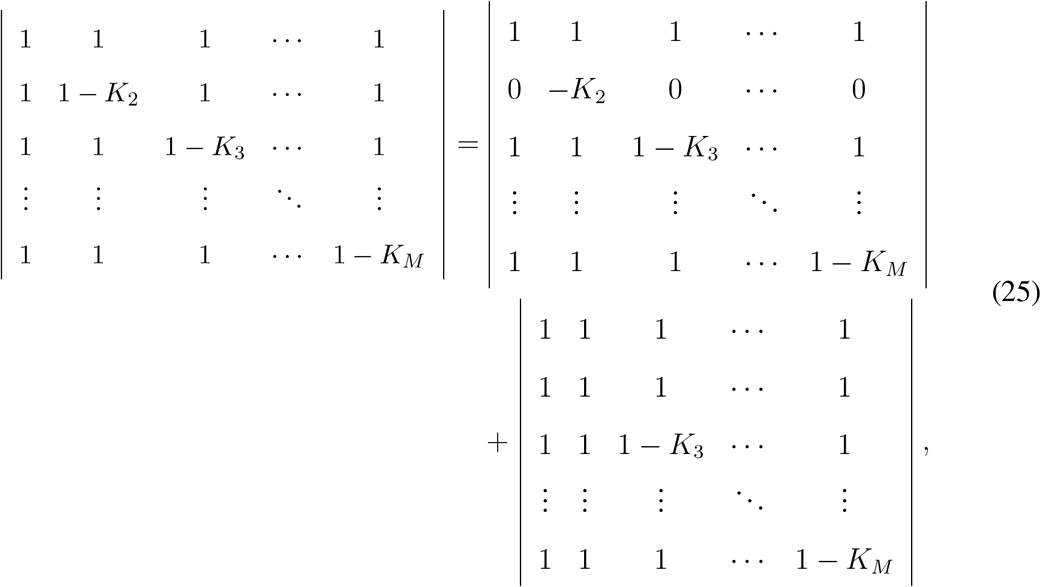

The second term in Eq. (25) is zero because the determinant has two identical columns, therefore only the first term remains. Continuing splitting the remaining rows of the first term of Eq. (25), we finally obtain

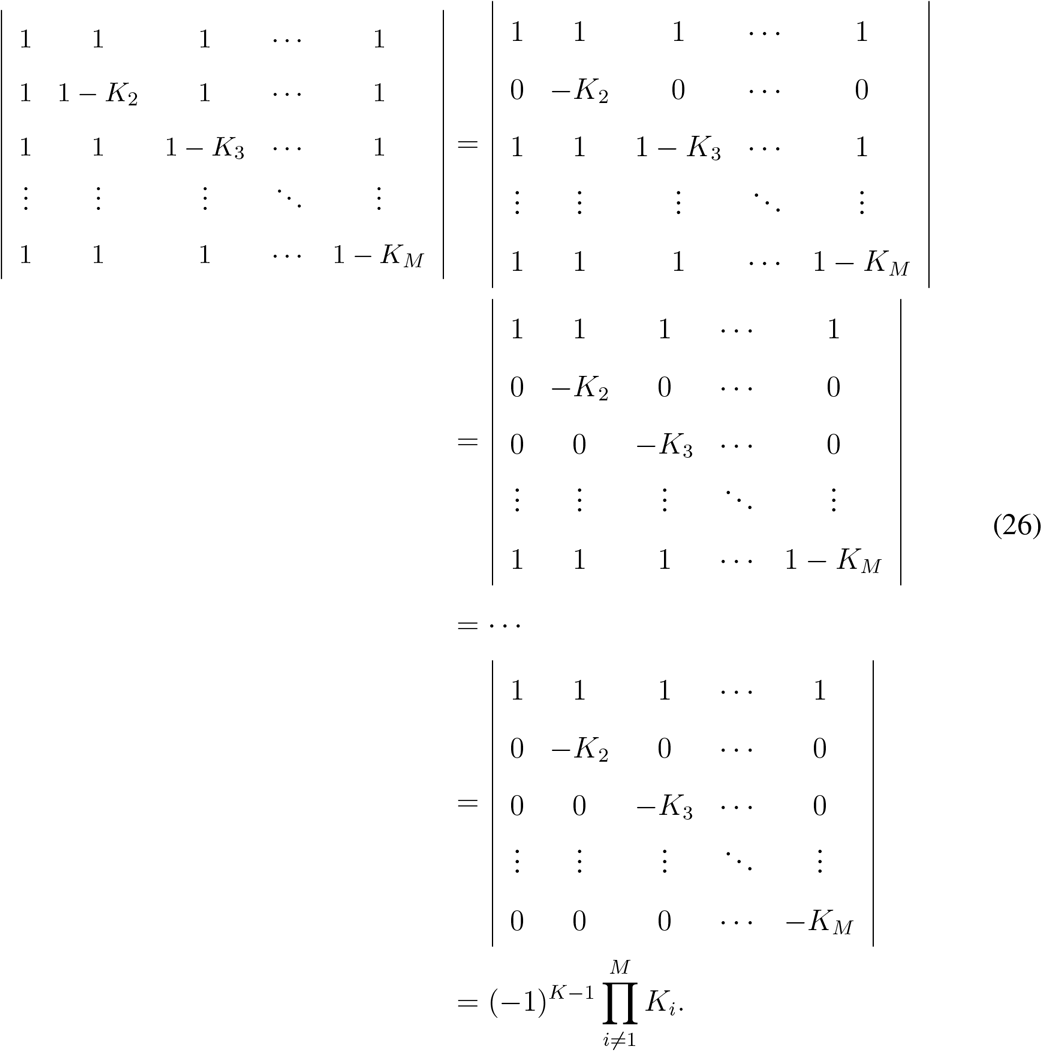

Now, we look back at the first term in Eq. (24), we split the second row

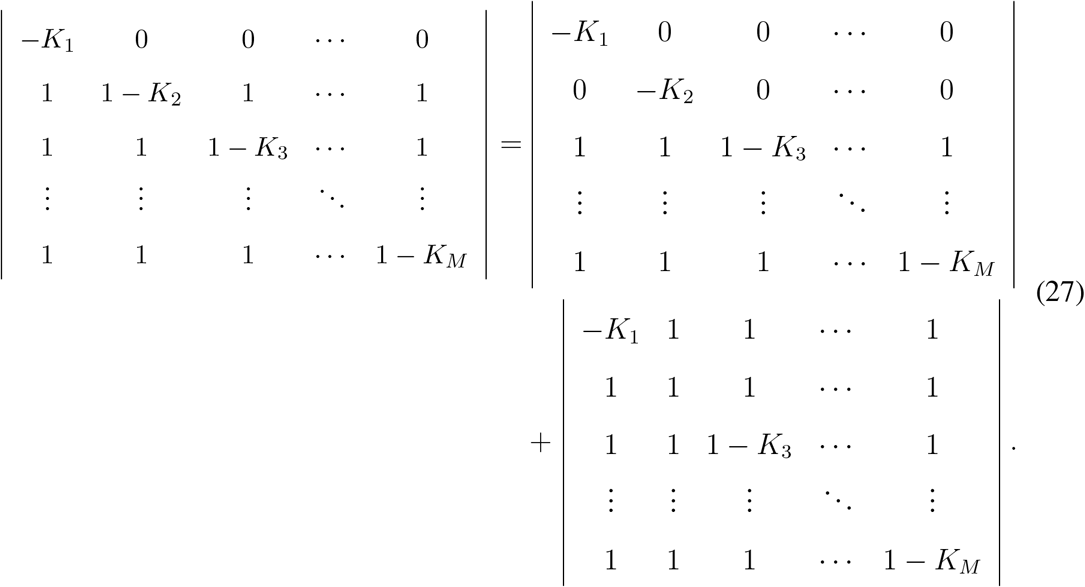

For the second term at the right hand side of Eq. (27), similar to Eq. (25) in the last step, we can work out that it equals 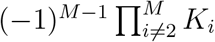. That means we can get 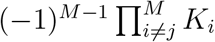 when split the *j*-th row. So we keep the same procedure to split the remaining rows of the first term in Eq. (27). After that, the initial determinant changes to

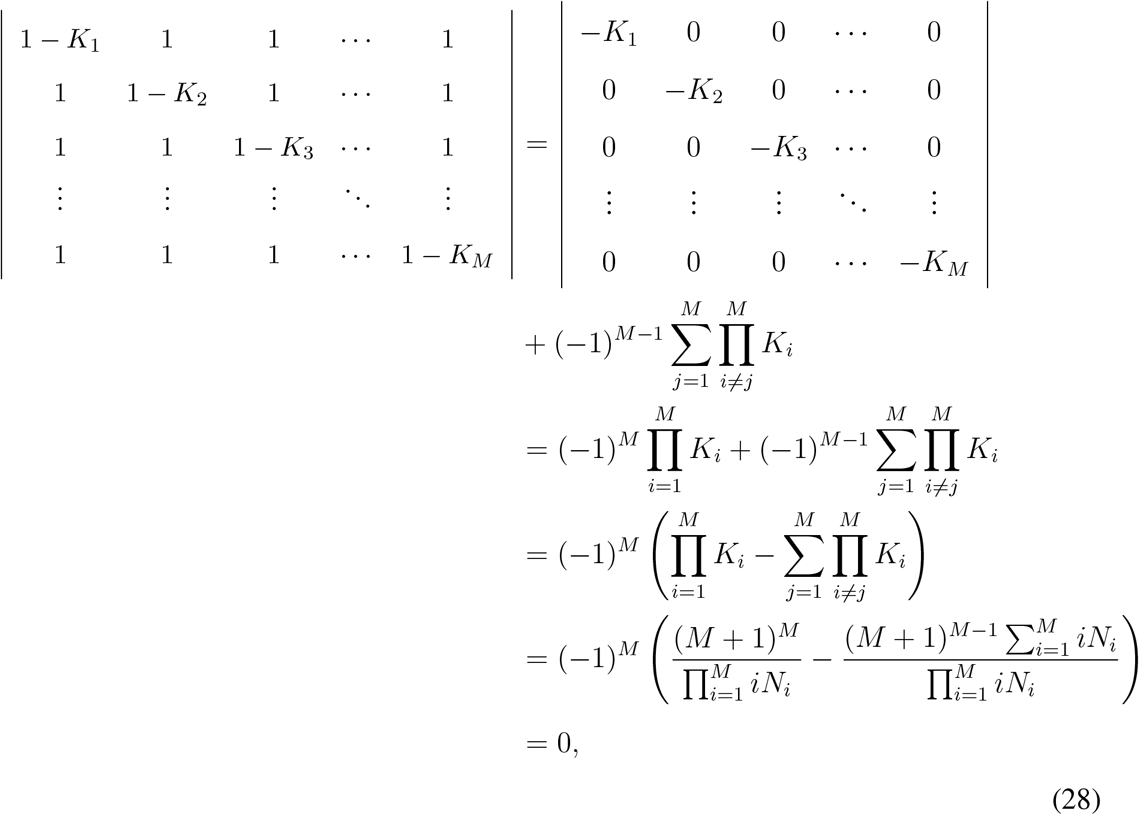

where we used 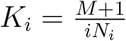 and 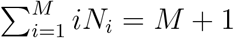 in the last two steps.

This proves that an arbitrary life cycle fragmenting at size *M* + 1 has the growth rate *λ* = 1, if 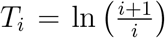 for any *i* ≤ *M*. This means that 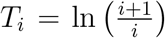 is a neutral fitness landscape for the scenario of homogeneous groups.

### A.3 Life cycles of homogeneous groups

In the absence of cells’ interactions, all cells are identical i.e. the cell type has no influence on groups. Essentially, all groups can be treated as homogeneous groups, in which only group sizes affect growth rate. In this case, groups have fixed developmental trajectories. For instance, the life cycle 1+1+1 has to go through the unique developmental trajectory: two successive divisions and then producing three single cells (see Fig. 5). In this unique developmental trajectory, only one initial type exist – independent cell, so *p*(*τ*) = 1 and **N**(*τ*) = 3.

**Figure 5:**
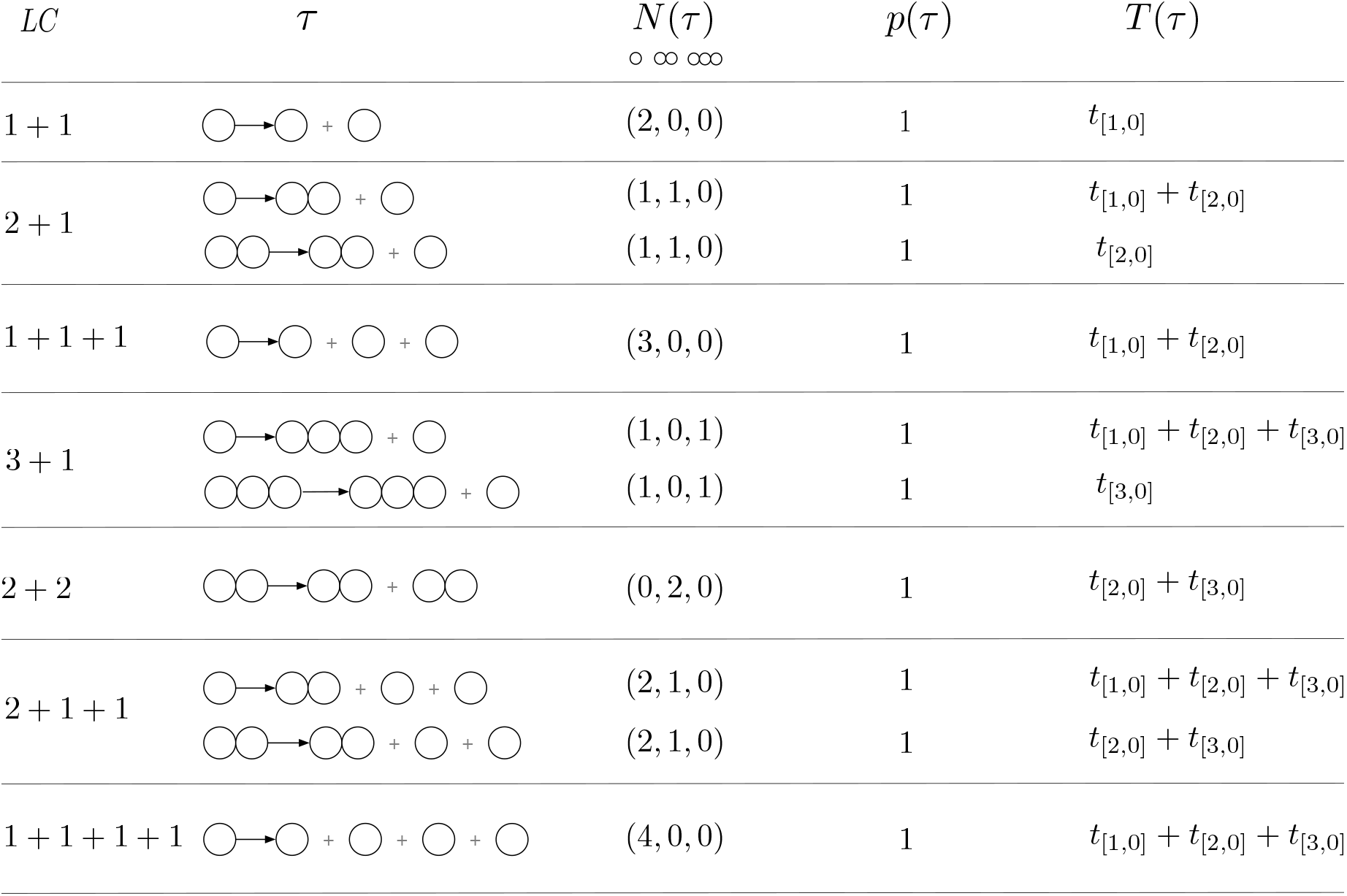
Homogeneous groups have deterministic developmental trajectories for each type offspring group, i.e. *p*(*τ*) = 1.

First, we investigate the simplest scenario, where the maximal size of the group was limited to two cells. There are three life cycles in total in this case: 1+1, 2+1 and 1+1+1. The matrices *Q* corresponding to these life cycles are

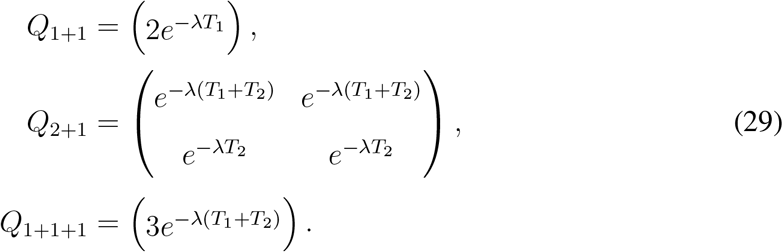

According to Eq. (4), the growth rate of each life cycle are given by the solutions of

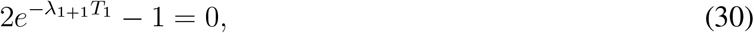

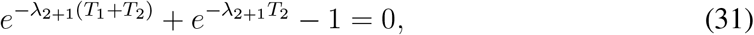

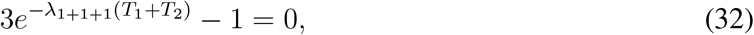

where *λ*_1+1_, *λ*_2+1_ and *λ*_1+1+1_ are the growth rate of 1+1, 2+1 and 1+1+1, respectively, see Fig. 6A. For small *T*_2_, the largest growth rate is achieved by 2+1 life cycle. In this case, bi-cellular groups produce offspring cells faster than independent cells. Consequently, the life cycle 2+1, which allows production of bi-cellular groups (unlike unicellular life cycle 1+1) and preserving one offspring group in the most productive bi-cellular state (unlike 1+1+1) is most successful in growth competition. In the opposite limit of large *T*_2_, the life cycle 1+1 leads to the largest population growth rate. In this case, independent cells are better off than bi-cellular groups. Thus, the best reproductive strategy is to avoid the growth to bi-cellular state, which can only be achieved with a single life cycle 1+1. In both situations of *T*_2_, the growth rate of 1+1+1 is always between that of 1+1 and 2+1.

**Figure 6:**
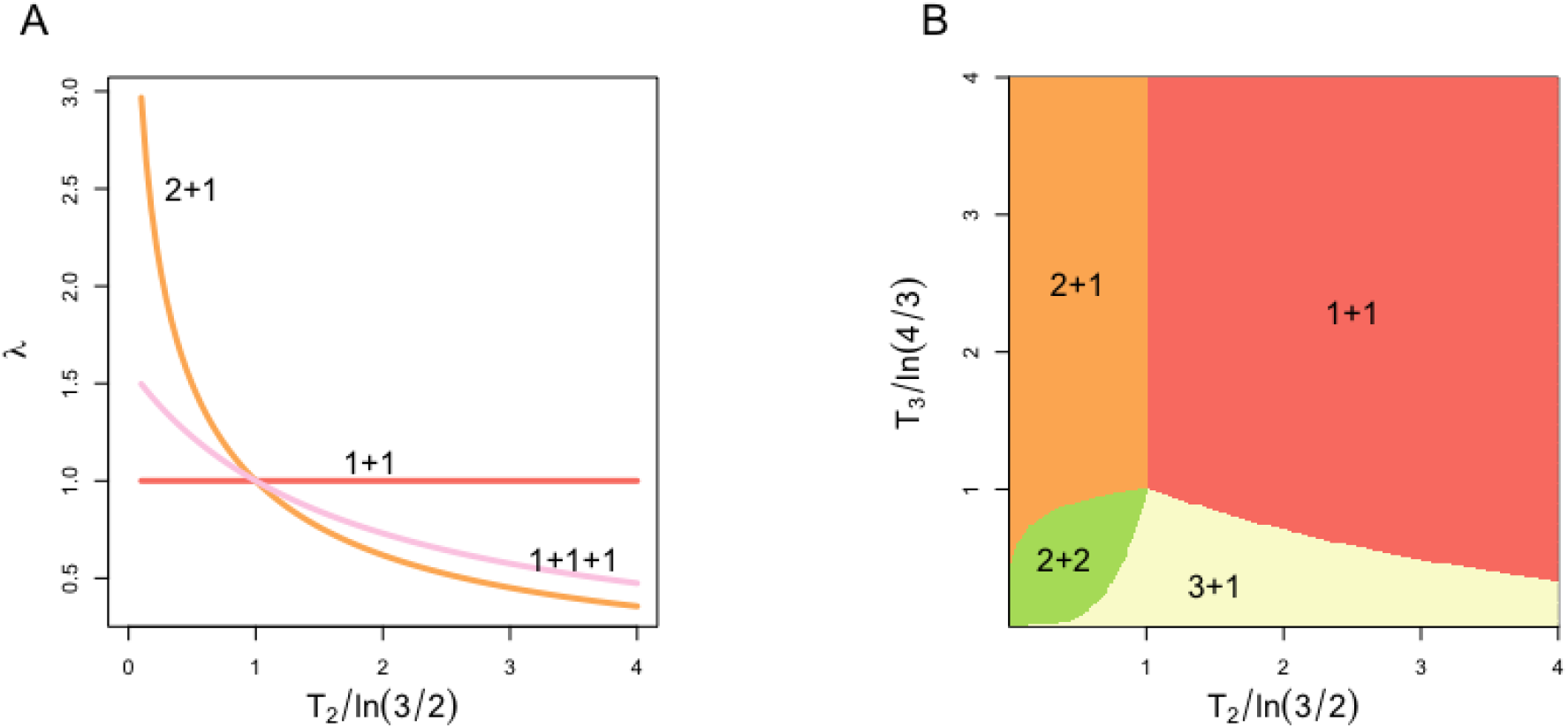
Growth rates and optimal life cycles in homogeneous groups on the condition of *n* ≤ 3 and *n* ≤ 4, respectively. **A**) describes the growth rates of life cycles when *n* ≤ 3 i.e. 1+1, 1+1+1 and 2+1. B) shows the optimal life cycle when *n* ≤ 4 with respect to *T*_2_ and *T*_3_. In both situations, *T_i_* is the size increment time and we set *T*_1_ = ln(2) for convenience.

For the next scenario, we increase the maximal size of the group to three cells. This allows four new life cycles: 3+1, 2+2, 2+1+1 and 1+1+1+1. Their growth rates are given by the solutions of

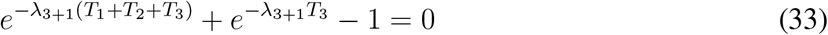

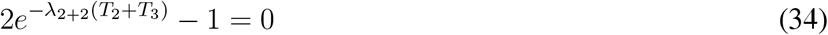

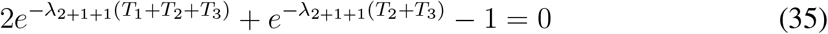

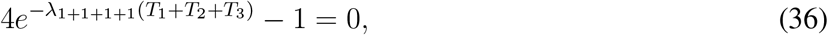

For large *T*_3_, the life cycles which do not produce slow-growing three-cellular groups have the highest growth rates. Therefore, for large *T*_3_, the optimal life cycles are the same as ones presented in the previous paragraph. For small *T*_3_, the life cycles capable of producing three-cellular groups gain an evolutionary advantage. Specifically, 2+2 achieves the maximum growth rate when both T2 and T3 are comparatively small. In this case, an independent cell is the least productive state, whereas 2+2 is the only life cycle not producing independent cells. Life cycle 3+1 leads to the largest growth rate if *T*_3_ is small but *T*_2_ is large. There, the three-cellular group stands out as the most productive state, and 3+1 is the only life cycle keeping it as one of its offspring groups. Similarly to the previous scenario, life cycles with more than two offspring: 1+1+1, 2+1+1, 1+1+1+1, are never optimal. An important exception to this is the point 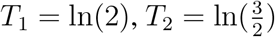 and 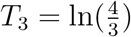, where all seven life cycles lead to the same growth rate (*λ* = 1).

Previously, we considered another model of life cycles evolution [Pichugin et al., 2017]. There, the growth of groups from size *i* to size *i* + 1 occurs spontaneously with rate *ib_i_*. Therefore, in that model, the time between cell divisions varies between groups of the same size, in contrast to the scenario considered here, where this time is always equal to *T_i_*. Despite the differences between two models, they both share a number of findings: existence of the neutral point, only binary fragmentation is evolutionarily optimal, same optimal life cycles in the limit cases. Therefore, these features, are independent from the model design.

### A.4 Calculation of growth rates *λ* for life cycles of heterogeneous groups

To show how our approach can can be used in the case of heterogeneous groups, consider the simplest unicellular life cycle 1+1. There are two types of offspring possible: independent *A* and *B* cells, so the matrix *Q* has dimensions 2 by 2. When a cell divides into two, three outcomes are possible: no cell, one cell, or both daughter cells change the phenotype. Since the developmental trajectory ends after the first division, there are only six developmental trajectories possible for this life cycle, see Fig. 7.

**Figure 7:**
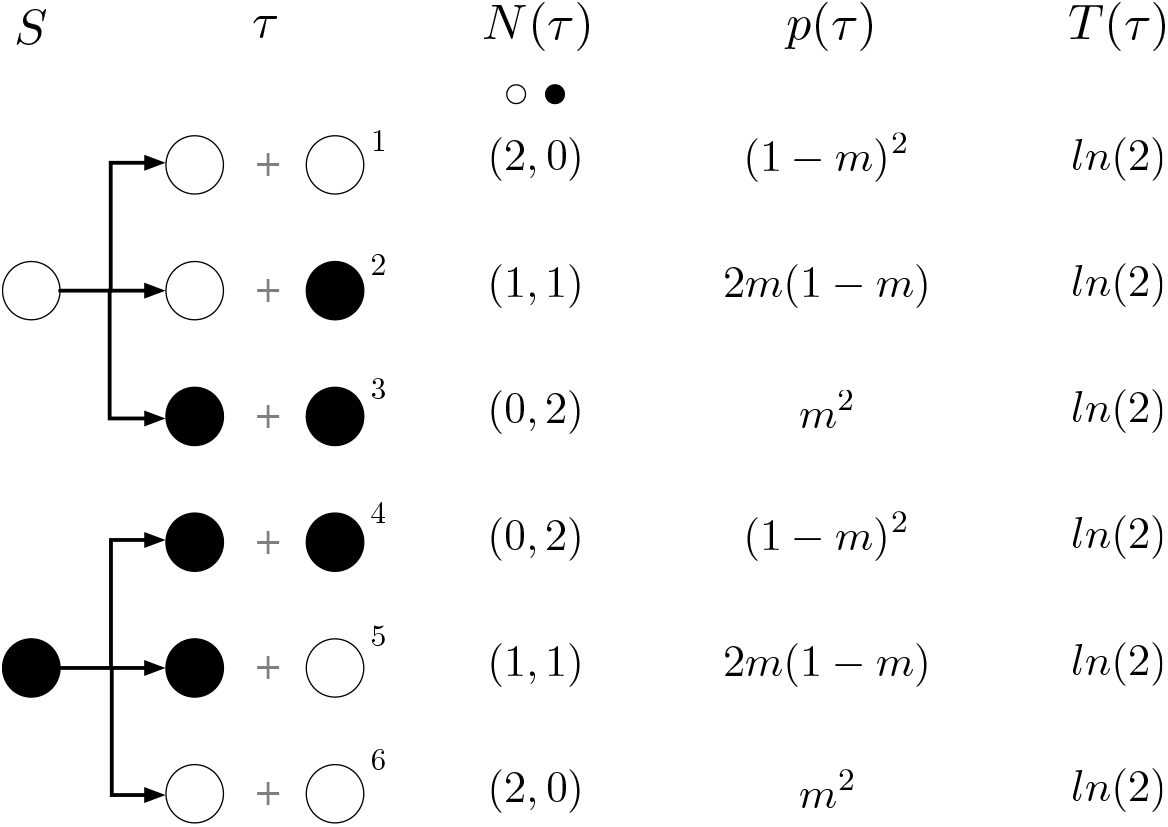
The full set of developmental trajectories in the life cycle 1+1. Here, the white and black circles denote *A* type cell and *B* type cell respectively.

To construct the matrix *Q*, we need to obtain the distribution of offspring (*N_i_*), the probability of realization (*p*) and total developmental time (*T*) for each trajectory. Offspring distributions are apparent from Fig. 7. The probability of each trajectory can be directly computed from the phenotype switch probability m. The developmental time is *T* = *T*_1_ = ln(2) for each trajectory here. Therefore, the elements of *Q* are given by

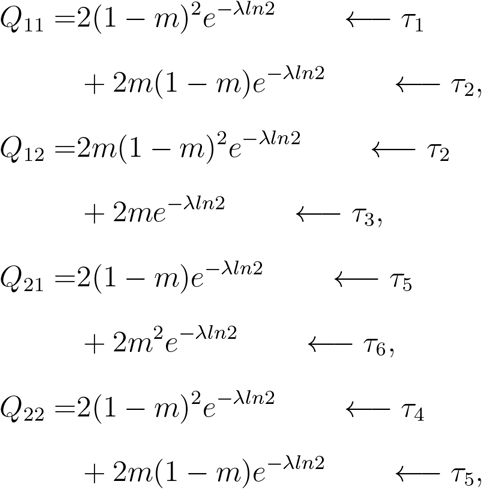

where arrows indicate the index of the developmental trajectory contributing a given term. Solution of the Eq. (18) leads to *λ*_1+1_ = 1 in the life cycle 1+1.

Next, consider the life cycle 1+1+1. There are still two types of offspring possible: independent *A* and *B* cells, such that the matrix *Q* has dimensions 2 by 2. However, the life cycle requires two divisions to complete, so the number of possible developmental trajectories is increased to 20. Also, interactions play a role during the second division, so the probabilities *p* and developmental times *T* are more complicated, see Fig. 8.

**Figure 8:**
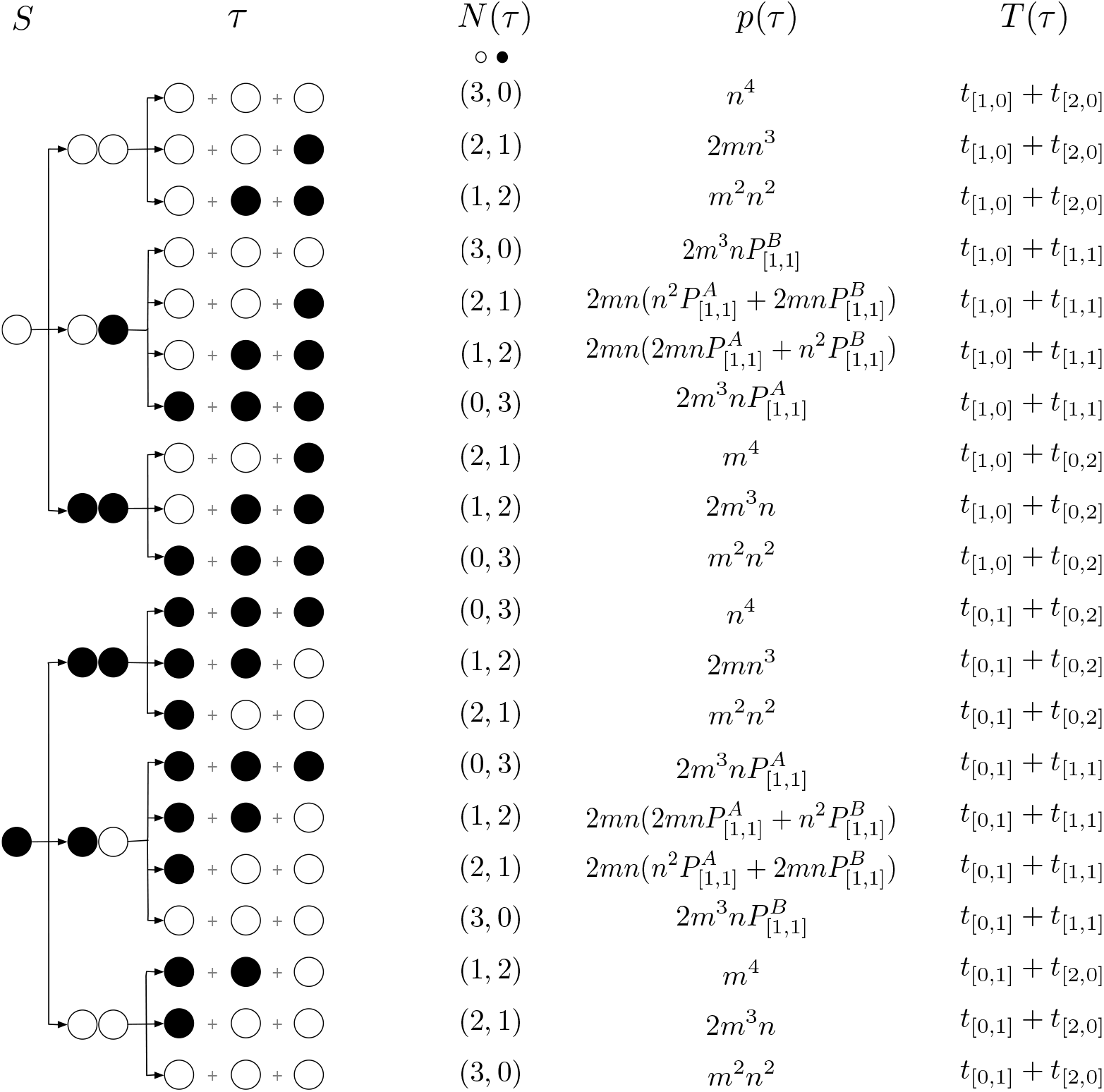
The full set of developmental trajectories in the life cycle 1+1+1. White circles represent *A* type cells and black circles represent *B* type cells. For simplicity of notation, we use *n* = 1 − *m, t*_[*i,j*]_ is the time before the next cell division for a complex with *i A* type cells and *j B* type cells to divide; and 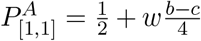 and 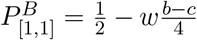, see details in the model section of the main text.

The elements of matrix *Q* are given by

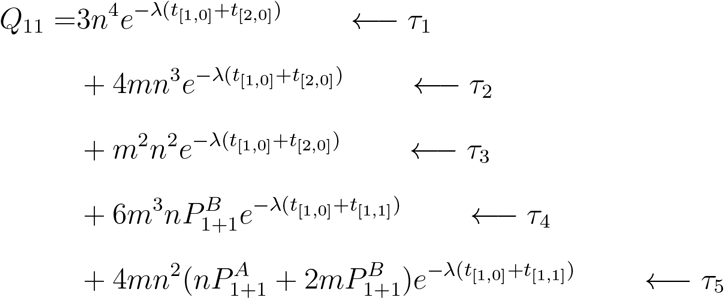

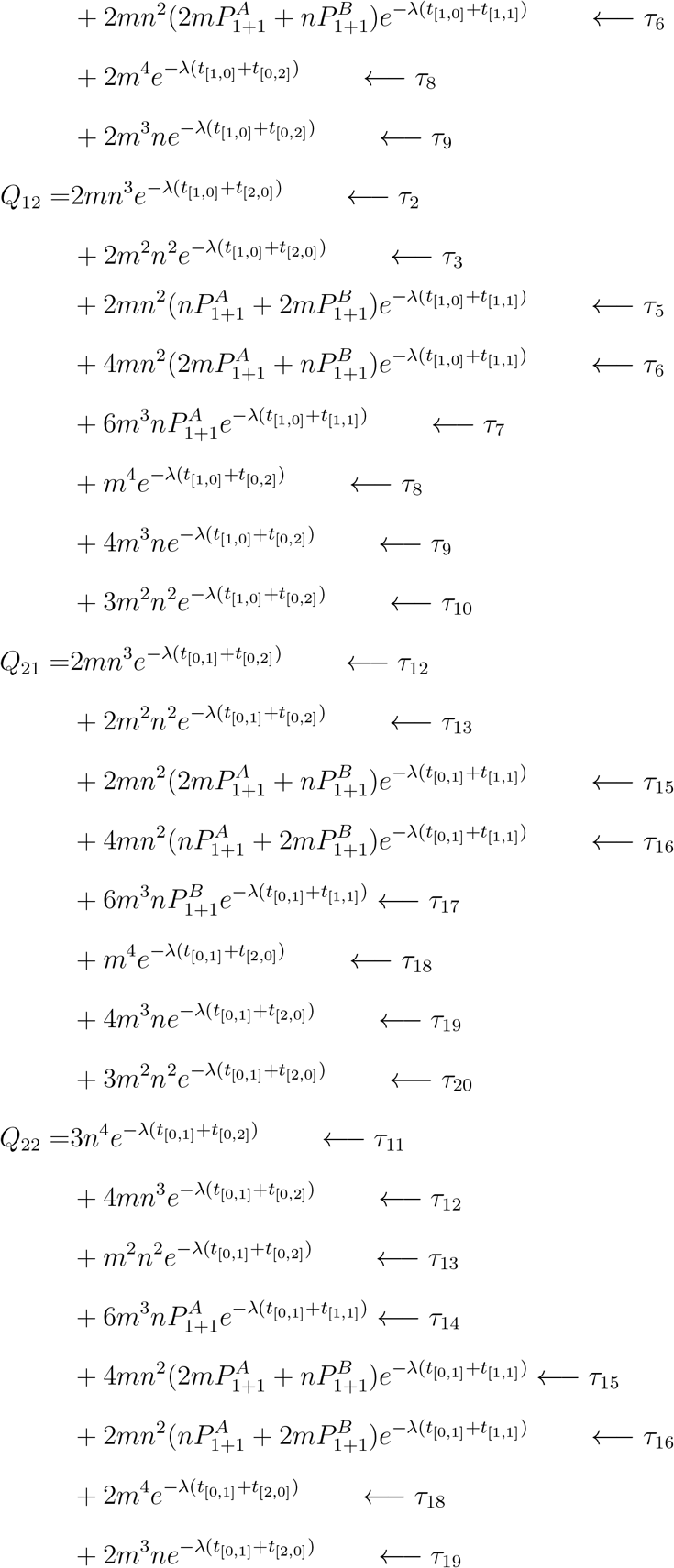

Here, 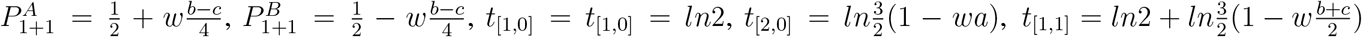 and 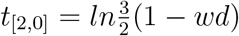. Arrows indicate the contributions of each developmental trajectory to *Q_ij_*.

The solution of Eq. (18) for life cycle 1+1+1 yields

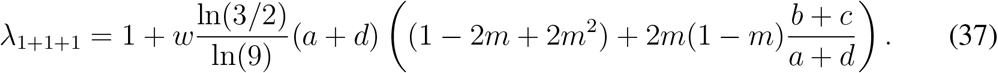

Our final example is the life cycle 2+1, where groups grow to three cells and fragment into a bi-cellular group and an independent cell. Here, five offspring types are possible: independent cells could be either *A* or *B* type and the bi-cellular group could have composition AA, AB, or BB, see Figs. 9,10. Therefore, *Q* is 5 × 5 matrix. There are 48 developmental programs possible and we refrain from showing here how elements of matrix *Q* are constructed in this case. The solution of the Eq. (18) for life cycle 2+1 yields

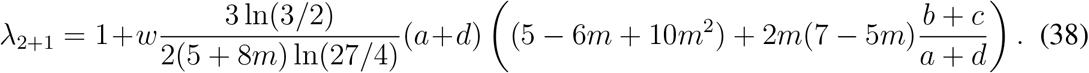

**Figure 9:**
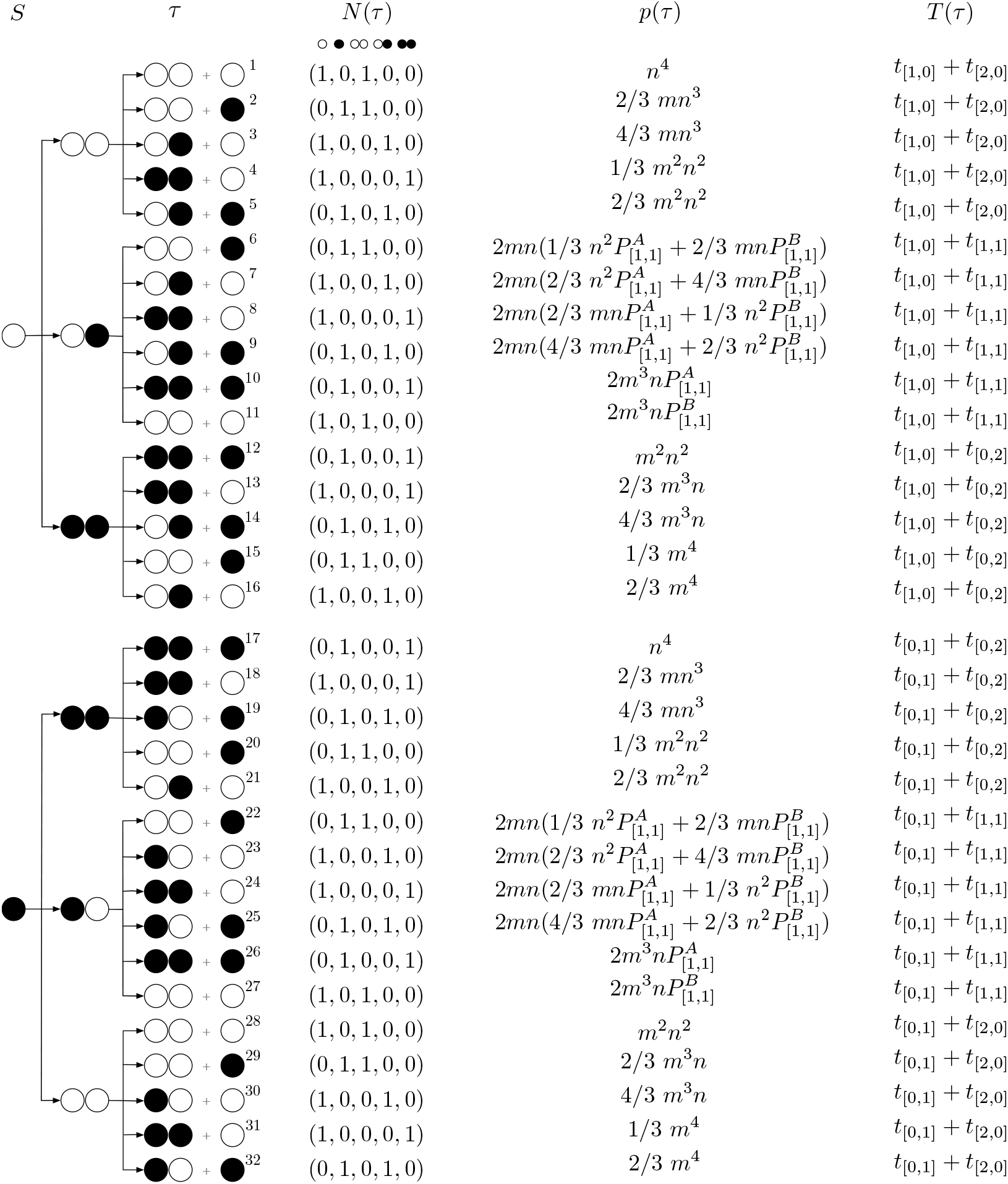
The developmental trajectories of the single A-cell and B-cell in the life cycle 2+1. White circles represent *A* type cells and black circles represent *B* type cells. For simplicity of notation, we use: *n* = 1 − *m*; *t*_[*i,j*]_ is the time before the next cell division for a complex with *i A* type cells and *j B* type cells to divide; and 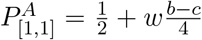 and 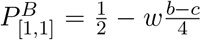, see details in the model section of the main text.

**Figure 10:**
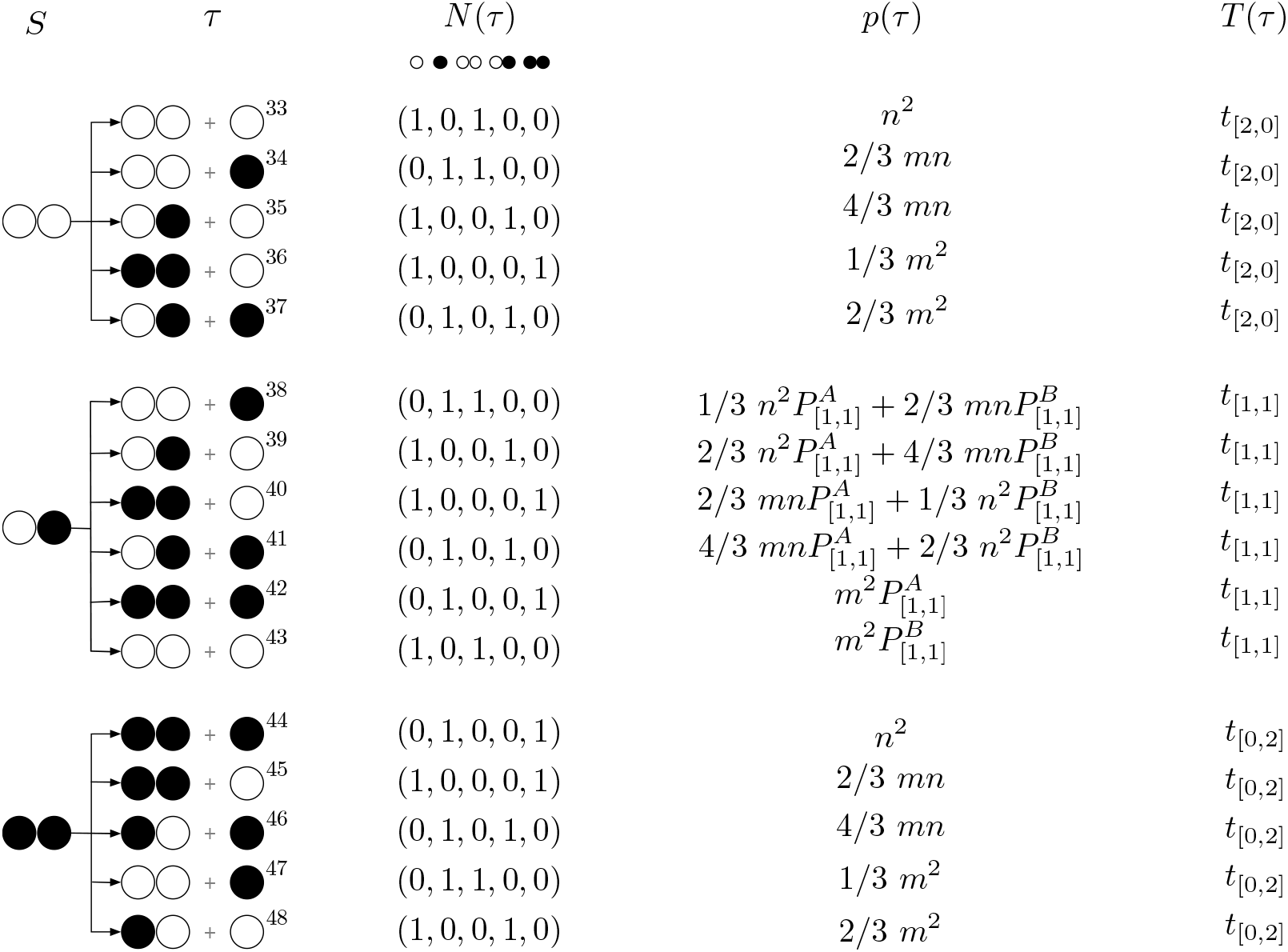
The developmental programs of bicellular newborn groups cells in the life cycle 2+1. White circles represent *A* type cells and black circles represent *B* type cells. For simplicity of notation, we use: *n* = 1 − *m*; *t*_[*i,j*]_ is the time before the next cell division for a complex with *i A* type cells and *j B* type cells to divide; and 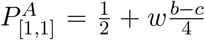 and 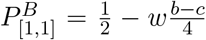, see details in the model section of the main text.

For the life cycles 1+1+1+1 and 2+2, we just list the growth rate *λ*

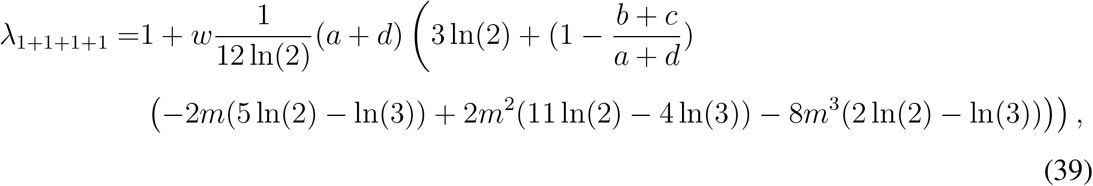

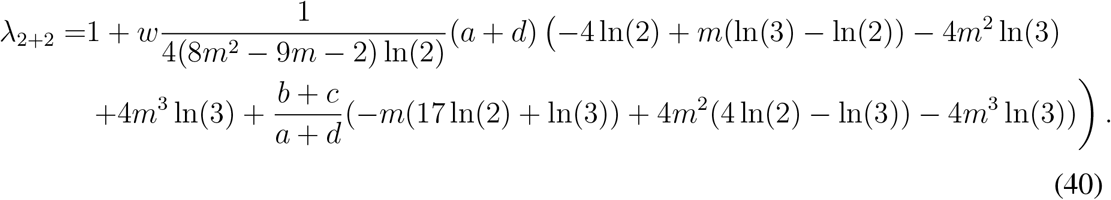

For more complex life cycles, the analytical expressions are too large to be meaningful by naked eye analysis. Therefore, we used a combination of analytical and numerical approaches. After the linearisation with respect to *w*, the growth rate for any life cycle has the form

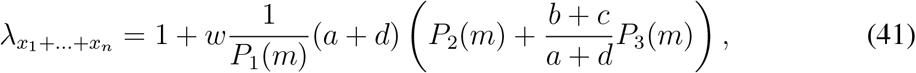

where *P*_1_(*m*), *P*_2_(*m*), *P*_3_(*m*) are some polynomials of m of the finite power. We obtained exact expressions for these polynomials using the symbolic algebra software. However, the tracking of all developmental programs means that the computation load grows exponentially with the maximal group size *M*. For life cycles, such as 3+2+1+1, computation of the polynomials required an extraordinary amount of RAM (> 70 Gb) and the outcome is neither human-tractable nor even printable. This memory constraints is the factor, limiting the maximal group size considered to *M* = 7. Therefore, in our study, we only stored the numerical values of the coefficients of *m* in *P*_1_(*m*), *P*_2_(*m*), *P*_3_(*m*) and used them to compute *λ*. With this approach, we are able to compute numerical values of *λ* with very high accuracy, even if traditional closed form solutions are unavailable.

### A.5 Profiles of growth rates of the life cycles

The growth rate is determined by three parameters: *ψ, ϕ* and *m*. The greatest diversity of evolutionarily optimal life cycles is observed at *ψ* > 0 and small *m*, see Fig. 4. In this appendix, we present profiles of growth rates at different conditions. For *ψ* > 0 and large m, only one multiple fission life cycle, 1+1+1, is evolutionary optimal, see Fig. 11A. Its area of optimality is located between unicellularity (1+1) at large negative *ϕ* and binary fragmentation with multicellular propagules (2+2 and 4+3) at large positive ϕ. Considering the dependence of growth rate from the phenotype switching probability *m*, we found that at *ϕ* ≫ 1, growth rate profiles are concave functions of *m*, see Fig. 11B. Growth rates of most life cycles are generally bound between binary fragmentation with multicellular propagules (such as 2+2 and 4+3) and multiple fragmentation with unicellular propagules (such as 1+1+1 and 1+1+1+1). For *ϕ* ≪ −1, the pattern is very similar, with an exception, that growth rate profiles are convex, instead of concave, and the hierarchy of life cycles is reversed, see Fig. 11C. This leads to the great diversity of evolutionary optimal life cycles at *ϕ* < 0 and small *m* (including also transitional life cycles 2+1 and 2+1+1, as well as binary fragmentation 2+2), see Fig. 11D.

**Figure 11:**
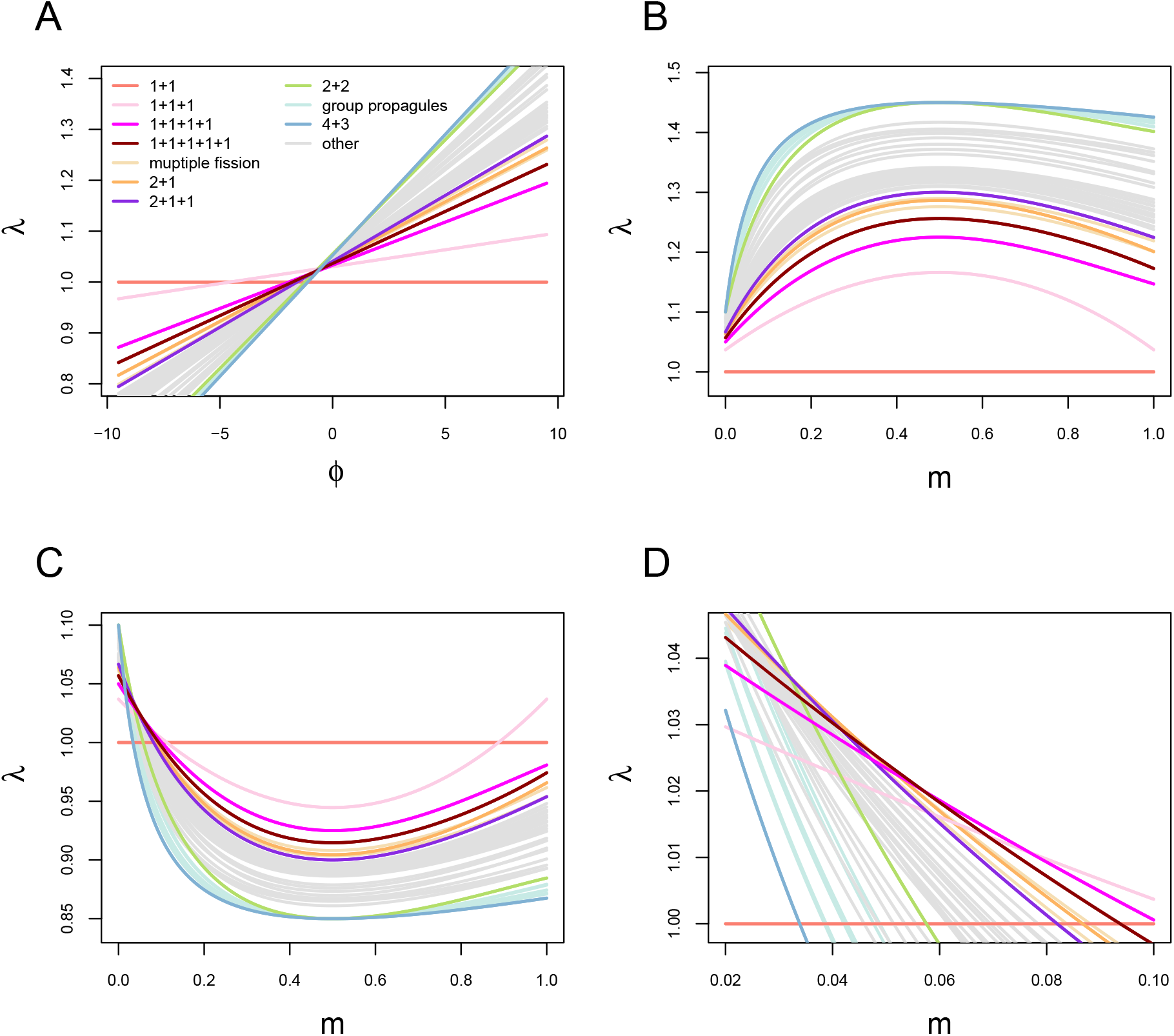
Multiple life cycles are optimal for *ψ* > 0. **A** Growth rates of all considered life cycles as function of *ϕ* at *m* = 0.9 (cf. Fig. 4A for *m* = 0.06). **B** Growth rates of all considered life cycles as function of *m* at *ϕ* = 8. **C** Growth rates of all considered life cycles as function of *m* at *ϕ* = −4. **D** Detailed view of the panel C in the range of small *m* showing that large number of evolutionary optimal life cycles at different *m*.

At the negative *ψ*, only two life cycles were found to be optimal, see Fig 12. The shape of individual growth rate profiles remain similar to the case of positive *ϕ* but the relative position changes significantly. Thus, the spectrum of observed life cycles is much less diverse.

**Figure 12:**
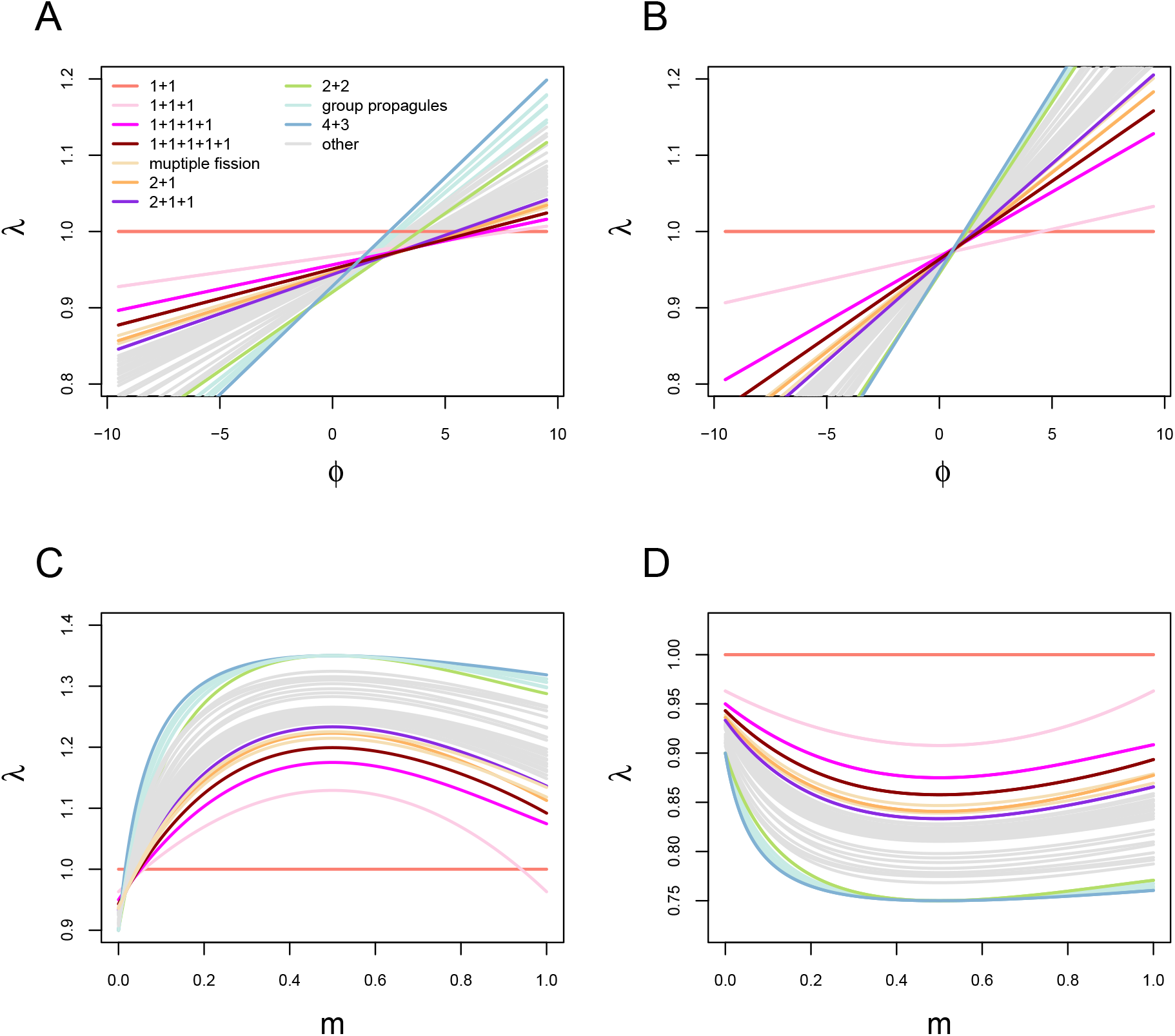
Only two life cycles are optimal for *ψ* < 0. **A** Growth rates of all considered life cycles as function of *ϕ* at *m* = 0.06 (cf. Fig. 4A for *ψ* > 0). **B** Growth rates of all considered life cycles as function of *ϕ* at *m* = 0.9 (cf. Fig. 11A for *ψ* > 0). **C** Growth rates of all considered life cycles as function of *m* at *ϕ* = 8 (cf. Fig. 11B for *ψ* > 0). **D** Growth rates of all considered life cycles as function of *m* at *ϕ* = −4 (cf. Fig. 11C for *ψ* > 0).

## References

J.T. Bonner. The Cellular Slime Molds. Princeton University Press, Princeton, NJ, 1959.

H. Brandt, C. Hauert, and K. Sigmund. Punishing and abstaining for public goods. Proceedings of the National Academy of Sciences USA, 103(2):495–497, 2006.

Dennis Claessen, Daniel E. Rozen, Oscar P. Kuipers, Lotte Sogaard-Andersen, and Gilles P. van Wezel. Bacterial solutions to multicellularity: a tale of biofilms, filaments and fruiting bodies. Nat Rev Micro, 12(2):115–124, 2014.

G.A. Cooper and S. A. West. Division of labour and the evolution of extreme specialization. Nature ecology & evolution, 2018.

S. De Monte and P. B. Rainey. Nascent multicellular life and the emergence of individuality. Journal of biosciences, 39(2):237 – 248, 2014.

Andre M De Roos. Demographic analysis of continuous-time life-history models. Ecology Letters, 11(1):1–15, 2008.

E. Flores and A. Herrero. Compartmentalized function through cell differentiation in filamentous cyanobacteria. Nature Reviews Microbiology, 8(1):39, 2010.

J. H. Fowler. Altruistic punishment and the origin of cooperation. Proceedings of the National Academy of Sciences USA, 102(19):7047–7049, 2005.

J. García and A. Traulsen. Leaving the loners alone: Evolution of cooperation in the presence of antisocial punishment. Journal of Theoretical Biology, 307:168–173, 2012.

Thomas Garcia, Leonardo Gregory Brunnet, and Silvia De Monte. Differential adhesion between moving particles as a mechanism for the evolution of social groups. PLoS Comput Biol, 10(2):e1003482, 02 2014.

Thomas Garcia, Guilhem Doulcier, and Silvia De Monte. The evolution of adhesiveness as a social adaptation. eLife, 4:e08595, 2015. doi: 10.7554/eLife.08595.

S. Gavrilets. Rapid transition towards the division of labor via evolution of developmental plasticity. PLoS Computational Biology, 6(6):e1000805, 2010.

Peter Godfrey-Smith. Darwinian Populations and Natural Selection. Oxford University Press, 2009.

R. K. Grosberg and J. E. Strassmann. The evolution of multicellularity: A minor major transition? Annual Review of Ecology, Evolution, and Systematics, 38:621–54, 2007.

K. Hammerschmidt, C. J. Rose, B. Kerr, and P. B. Rainey. Life cycles, fitness decoupling and the evolution of multicellularity. Nature, 515(7525):75–79, 2014.

G. Hardin. The tragedy of the commons. Science, 162:1243–1248, 1968.

C. Hauert, S. De Monte, J. Hofbauer, and K. Sigmund. Volunteering as red queen mechanism for cooperation in public goods games. Science, 296:1129–1132, 2002.

C. Hauert, A. Traulsen, H. Brandt, M. A. Nowak, and K. Sigmund. Via freedom to coercion: the emergence of costly punishment. Science, 316:1905–1907, 2007.

C. Hilbe, M. A. Nowak, and K. Sigmund. The evolution of extortion in iterated prisoner’s dilemma games. Proceedings of the National Academy of Sciences USA, 110:6913–6918, 2013.

J. Hofbauer and K. Sigmund. Evolutionary Games and Population Dynamics. Cambridge University Press, Cambridge, UK, 1998.

Iaroslav Ispolatov, Martin Ackermann, and Michael Doebeli. Division of labour and the evolution of multicellularity. Proceedings of the Royal Society of London B: Biological Sciences, 279(1734):1768–1776, 2012.

K. Kaveh, C. Veller, and M. A. Nowak. Games of multicellularity. Journal of Theoretical Biology, 403:143 – 158, 2016.

A. H. Knoll. The early evolution of eukaryotes: a geological perspective. Science, 256(5057):622–627, 1992.

E. Libby, W. C. Ratcliff, M. Travisano, and B. Kerr. Geometry shapes evolution of early multicellularity. PLoS Computational Biology, 10(9):e1003803, 2014.

J. Maynard Smith. Evolution and the Theory of Games. Cambridge University Press, Cambridge, 1982.

R. E. Michod and D. Roze. Some aspects of reproductive mode and origin of multicellularity. Selection, 1(1-3):97 – 110, 2001.

Richard E Michod. Cooperation and conflict in the evolution of individuality. i. multilevel selection of the organism. The American Naturalist, 149:607–645, 1997.

M. A. Nowak. Five rules for the evolution of cooperation. Science, 314:1560–1563, 2006a.

M. A. Nowak. Evolutionary dynamics. Harvard University Press, Cambridge MA, 2006b.

M. A. Nowak and K. Sigmund. Evolutionary dynamics of biological games. Science, 303: 793–799, 2004.

S. Okasha. Evolution and the levels of selection. Oxford Univ. Press, 2006.

J. M. Pacheco, F. C. Santos, M. O. Souza, and B. Skyrms. Evolutionary dynamics of collective action in n-person stag hunt dilemmas. Proceedings of the Royal Society B, 276: 315–321, 2009.

Y. Pichugin and A. Traulsen. Competition of multicellular life cycles under costly reproduction. bioRxiv, in press 2018.

Y. Pichugin, J. Peña, P. Rainey, and A. Traulsen. Fragmentation modes and the evolution of lifecycles. PLoS Computational Biology, 13(11):e1005860, 2017.

P. B. Rainey and B. Kerr. Cheats as first propagules: a new hypothesis for the evolution of individuality during the transition from single cells to multicellularity. BioEssays, 32(10): 872–880, 2010.

P. B. Rainey and K. Rainey. Evolution of cooperation and conflict in experimental bacterial populations. Nature, 425(6953):72–74, 2003.

Armin Rashidi, Deborah E. Shelton, and Richard E. Michod. A darwinian approach to the origin of life cycles with group properties. Theoretical Population Biology, 102:76 – 84, 2015.

W. C Ratcliff, R. F Denison, M Borrello, and M Travisano. Experimental evolution of multi-cellularity. Proceedings of the National Academy of Sciences USA, pages 1–6, Jan 2012.

W. C. Ratcliff, M. D. Herron, K. Howell, J. T. Pentz, F. Rosenzweig, and M. Travisano. Experimental evolution of an altering uni- and multicellular life cycle in chlamydomonas reinhardtii. Nature Communications, 4(2742), November 2013a.

W. C. Ratcliff, J. T. Pentz, and M. Travisano. Tempo and mode of multicellular adaptation in experimentally evolved Saccharomyces cerevisiae. Evolution, 67(6):1573–1581, 2013b.

João F Matias Rodrigues, Daniel J Rankin, Valentina Rossetti, Andreas Wagner, and Homay-oun C Bagheri. Correction: Differences in Cell Division Rates Drive the Evolution of Terminal Differentiation in Microbes. PLoS computational biology, 8(5), 2012.

Valentina Rossetti, Manuela Filippini, Miroslav Svercel, A. D. Barbour, and Homayoun C. Bagheri. Emergent multicellular life cycles in filamentous bacteria owing to density-dependent population dynamics. Journal of The Royal Society Interface, 8(65):1772–1784, 2011. doi: 10.1098/rsif.2011.0102.

D. Roze and R. E. Michod. Mutation, multilevel selection, and the evolution of propagule size during the origin of multicellularity.”. The American Naturalist, 158(6):638 – 654, 2001.

C. E. Tarnita, C. H. Taubes, and M. A. Nowak. Evolutionary construction by staying together and coming together. Journal of Theoretical Biology, 320(0):10–22, 2013.

A. Tomitani, A. H. Knoll, C.M. Cavanaugh, and T. Ohno. The evolutionary diversification of cyanobacteria: molecular–phylogenetic and paleontological perspectives. Proceedings of the National Academy of Sciences, 103(14):5442–5447, 2006.

A. Traulsen, C. Hauert, H. De Silva, M. A Nowak, and K. Sigmund. Exploration dynamics in evolutionary games. Proceedings of the National Academy of Sciences USA, 106:709–712, 2009.

J. van Gestel and C. E. Tarnita. On the origin of biological construction, with a focus on multicellularity. Proceedings of the National Academy of Sciences, 114(42):11018 – 11026, 2017.

M. van Veelen. Group selection, kin selection, altruism and cooperation: When inclusive fitness is right and when it can be wrong. Journal of Theoretical Biology, 259:589–600, 2009.

J. W. Weibull. Evolutionary Game Theory. MIT Press, Cambridge, 1995.

